# Inflammatory Stromal Aging in Ulcerative Colitis and Colitis-Associated Cancer

**DOI:** 10.64898/2026.05.18.725891

**Authors:** Khalil Almotah, Uyen Tran, Haylee Gilbert, R. Allen Schweickart, Robert C. Fisher, Yaroslav Bisikalo, Manal Ali, Munir Buhaya, Mingyuan Cheng, Michael Cruise, Zhikai Chi, Samaneh Kamali Sarvestani, Emina H. Huang, Oliver Wessely

**Affiliations:** Dept. Cardiovascular & Metabolic Sciences, Cleveland Clinic Research, Cleveland Clinic, 9500 Euclid Avenue, Cleveland OH 44195; Dept. Chemistry, College of Arts and Sciences, Cleveland State University, 2121 Euclid Avenue, Cleveland, OH 44115; Dept. Inflammation and Immunity, Cleveland Clinic Research, Cleveland Clinic, 9500 Euclid Avenue, Cleveland OH 44195; Dept. of Surgery, UT Southwestern Dallas, 5323 Harry Hines Boulevard, Dallas, Texas 75390; Simmons Comprehensive Cancer Center, University of Texas Southwestern, 5323 Harry Hines Boulevard, Dallas, TX 75390; Dept. of Pathology, Cleveland Clinic Lerner College of Medicine, Cleveland Clinic, 9500 Euclid Avenue, Cleveland OH 44195; Dept. of Pathology, University of Texas Southwestern Medical Center, 5323 Harry Hines Boulevard, Dallas, TX 75390; Dept. of Molecular Medicine, Cleveland Clinic Lerner College of Medicine, Case Western Reserve University, 10900 Euclid Avenue, Cleveland, OH 44106

**Keywords:** Colitis-associated cancer, CXCL8, fibroblasts, iPSC reprogramming, NF-κB signaling, Senescence-associated secretory phenotype (SASP), ulcerative colitis

## Abstract

Ulcerative colitis is a chronic inflammatory bowel disease that can progress from dysplasia to cancer. Inflammatory responses are critical drivers in this process, typically triggered by epithelial lesions and the ensuing infiltration of microbiota into the interstitial layer. Here, we focus on the pro-inflammatory state of the interstitial fibroblasts, which promotes immune infiltration and augments disease progression. The study aims to provide a mechanistic link how fibroblasts of the colitis-associated microenvironment integrate inflammatory signals, microbial infiltration and cellular memory. To this end, we investigated a large number of primary colon fibroblasts obtained from normal, colitis and colon cancer samples using a range of *in vitro* approaches and an *in vivo* co-inoculation cancer model. mRNA sequencing analysis identified that the disease-associated fibroblasts are exhibit a cellular inflammatory status, which involves the injury-induced senescence pathway. Using CXCL8, a potent chemokine upregulated in colitis and cancer colon fibroblasts, as a paradigm, this inflammatory status is triggered by the activation of the NFκB signaling via immune-derived cytokines (TNFα, IL-1β), bacterial signals (LPS) and the microbiome itself using mycoplasma as a paradigm. Finally, iPSC reprogramming studies indicate that fibroblasts from ulcerative colitis retain an epigenetic memory that sustains elevated CXCL8 expression. Together, our findings demonstrate that the senescence associated secretory phenotype of colon fibroblasts is a robust indicator for inflammation-driven colon tumorigenesis.

## BACKGROUND

Inflammatory Bowel Disease (IBD), which includes Ulcerative Colitis (UC) and Crohn’s Disease (CD), is characterized by chronic relapsing inflammation of the gastrointestinal tract resulting from an abnormal immune response. This dysregulation leads to cycles of remission and recurrence that can span decades and affect individuals of all ages [1–4]. In patients with long-standing UC, unresolved chronic inflammation increases the risk for the development of colitis-associated cancer (CAC) [5–7]. The development of dysplasia in UC and the disease progression to cancer has been linked to immune cell recruitment and ensuing chemokine- and cytokine-driven interactions with epithelial and stromal cells [8–10]. These changes may lead to molecular alterations and can trigger genetic mutations that drive the progression of UC towards CAC.

In the dysplastic tissue, ulcerative colitis-associated fibroblasts (UCAFs) and cancer-associated fibroblasts (CAFs) have been implicated in tumor initiation and progression [1, 11]. Fibroblasts in the colonic microenvironment can respond to signals from epithelial cells and inflammatory cues, adopting an activated state associated with pro-inflammatory and tumor-promoting functions [12]. UCAFs and CAFs influence the immune landscape in the colon by secreting growth factors, cytokines, and extracellular matrix components that can suppress anti-tumor immunity, thereby promoting tumor initiation and progression via cancer cell proliferation, invasion, and metastasis.[13–15] Furthermore, UCAFs and CAFs modulate the immune system within the colon by suppressing the immune microenvironment [16, 17], ultimately promoting tumor initiation, growth and survival. Recognition of UCAFs and CAFS as active contributors to tumorigenesis has stimulated interest in therapeutic strategies targeting the tumor microenvironment, including fibroblast-mediated signaling and stromal remodeling [18, 19].

Chronic inflammation can promote a senescence-associated secretory phenotype (SASP) in UCAFs and CAFs, resulting in the secretion of proinflammatory and pro-tumorigenic mediators [20, 21]. SASP factors include cytokines, chemokines, growth factors, and proteases that alter tissue homeostasis and influence cellular function, differentiation, and proliferation ultimately providing a microenvironment supportive for tumor growth [21]. Among the numerous proinflammatory secreted modulators within the SASP, the chemokine CXCL8 plays a crucial role in the inflammatory responses associated with UC.[22–24] Elevated levels of CXCL8 have been found in colon biopsies of UC and colon cancer patients, indicating a potential link between CXCL8 and the development of CAC.[23, 25] While CXCL8 is secreted by epithelial, immune, and endothelial cells,[26–29] recent data highlight a significant contribution from fibroblasts.[23] Thus, fibroblast-derived CXCL8 may play an important role in mediating tumor-stromal crosstalk in CAC.[30]

The present study focuses on the regulation of CXCL8 secretion in primary human colon fibroblasts as a paradigm for the fibroblast-epithelial and stromal-microbial crosstalk in the inflamed and neoplastic colon. We show that fibroblasts from human colitis patients exhibit an upregulated inflammatory phenotype compared to their normal colon healthy counterparts. The inflammatory signature is further aggravated upon transition to CAC. We demonstrate that CXCL8 expression is regulated by TNFα, IL-1β and LPS through NF-κB signaling. Additionally, mycoplasma infection as a microbial stimulus is capable of triggering similar responses, exemplifying the impact of the microbiome on stromal inflammation. This interaction is disease relevant *in vivo* as UCAFs augment tumor formation in mice, which is further augmented by TNFα. Finally, we show that the enhanced CXCL8 response of UCAFs over normal colon fibroblasts is retained through reprogramming into induced pluripotent stem cells (iPSCs) and differentiation into mesenchymal-derived fibroblasts, suggesting an epigenetically encoded proinflammatory phenotype. Together, our findings position fibroblast-derived CXCL8 as a robust marker and potential target in inflammation-driven colon tumorigenesis.

## MATERIALS AND METHODS

### Human Patient Material

Patients with ulcerative colitis or colitis-associated cancer were consented under IRB 13-1159 (Cleveland Clinic) or IRB 2021-0161 (University of Texas, Southwestern) for retrieval of surgically resected tissue samples. Demographics are presented in **Supplementary Table S1**. The human tissues were collected under pathologic observation, and normal colon tissues were sourced from locations at least 10 cm away from evident tumors. Pathological review was provided by Dr. Michael Cruise (Cleveland Clinic) and Dr. Zhikai Chi (University of Texas, Southwestern) (**Supplementary Table S1**).

### Primary Human Fibroblast Isolation

Surgically resected tissue samples were sterilized by dipping in dilute bleach for 30 seconds followed by treatment with antifungals and antimicrobials (#15240-062, Gibco) 3 times for 15 minutes each. Tissues were minced in 500-1000 μl DMEM containing 1 mg/ml collagenase Type 1A (#C9891, Sigma) in a round bottom 2 mL microfuge tube until the homogenate could pass through a 1000 μl pipette tip. Minced tissue was transferred to a 50 ml conical tube with an addition of 9 ml of collagenase-containing media, then placed in a shaking water bath at 37°C for 30 minutes. Collagenase activity was terminated by adding serum-containing DMEM media and filtering through a 40 µm strainer. The filtrate was cultured on tissue culture plates in DMEM supplemented with 10% FBS and 1% Pen/Strep.

Short-tandem repeat (STR) analyses to confirm the identity of the primary fibroblast/organoid isolates was performed upon genomic DNA extraction (DNEasy Blood and Tissue Kit, #69506, Qiagen) using Duke University (10 markers), Genetica (16 markers), or the McDermott genomics core at the University of Texas Southwestern (24 markers) (**Supplementary Table S2**). Concordance of >90% of the markers was defined as identity validity. For primary isolates, the cellular derivatives were compared to genomic DNA isolated from parental formalin-fixed paraffin-embedded sections (QIAamp DNA FFPE Tissue Kit, #56404, Qiagen).

The generation of the CT5-SV40-largeT-TERT ulcerative colitis-associated fibroblast line was described previously [30]. This isolate was established from an acute colitic patient at the University of Florida, then immortalized with the SV40-Large T antigen in a lentiviral construct (kind gift of Rosa Hwang, MD Anderson). The human colon fibroblast cell line (CRL1541) and colon cancer epithelial cell line (HCT 116) were obtained from ATCC, validated via STR analyses and tested for mycoplasma contamination prior to use.

### Vertebrate Animals

All animal procedures were approved by the Institutional Animal Care and Use Committee (IACUC) of the University of Texas Southwestern Medical Center and conducted in accordance with institutional guidelines and the ARRIVE guidelines for the care and use of laboratory animals (IACUC protocol #103065). A breeding colony of immunodeficient non-obese diabetic IL2 gamma-receptor null (NOD-SCID IL2 γ receptor null, here referred to as NSG) mice was established by the Simmons Comprehensive Cancer center at the University of Texas Southwestern. Experimental groups consisted of at least 5 mice between 7-9 weeks-old with a 1.3:1 F:M ratio. Mice were monitored every 2–3 days, and tumor volume was determined using digital calipers. Animals were euthanized prior to tumors reaching 2 cm in diameter, as required by IACUC policy. Euthanasia was performed using a lethal dose of ketamine–xylazine followed by cervical dislocation. No animals exceeded the humane endpoints outlined in the protocol.

### Statistics

All statistical analyses for the *in vitro* studies were conducted using GraphPad Prism (version 10). Statistical significance was defined as P < 0.05. In all cases a minimum of 3 biological and technical replicates were used.

## RESULTS

### Expression of CXCL8 in Primary Human Colitis-Associated Fibroblasts

To investigate changes in CXCL8 expression in human colon fibroblasts during progression towards colitis-associated cancer, we established mycoplasma-free primary fibroblast cultures from normal colon tissue, ulcerative colitis and colon cancer (including both CAC and sporadic colon cancer). In agreement with our previous reports,[30, 31] both ELISA and qRT-PCR analyses demonstrated upregulated levels of CXCL8 mRNA and protein in the UCAFs compared to normal fibroblasts (**Figure 1A,B**). As CXCL8 is part of the senescence associated secretory phenotype (SASP)[32] we next evaluated the expression of the canonical senescence markers including *CDKN2A/p16*, *TP53* and *CDKN1A/p21* mRNA levels (**Figure 1C-E**). *TP53* was significantly upregulated, with a trend toward increased *CDKN1A* expression in UCAFs, whereas *CDKN2A* levels remained unchanged. This pattern suggests that the UCAFs exhibit injury/stress-induced senescence compared to normal colon fibroblasts, rather than replicative senescence. Of note the cancer-associated fibroblasts (CAFs) trended in the same direction but showed a less pronounced response than the UCAFs.

**Figure 1:**
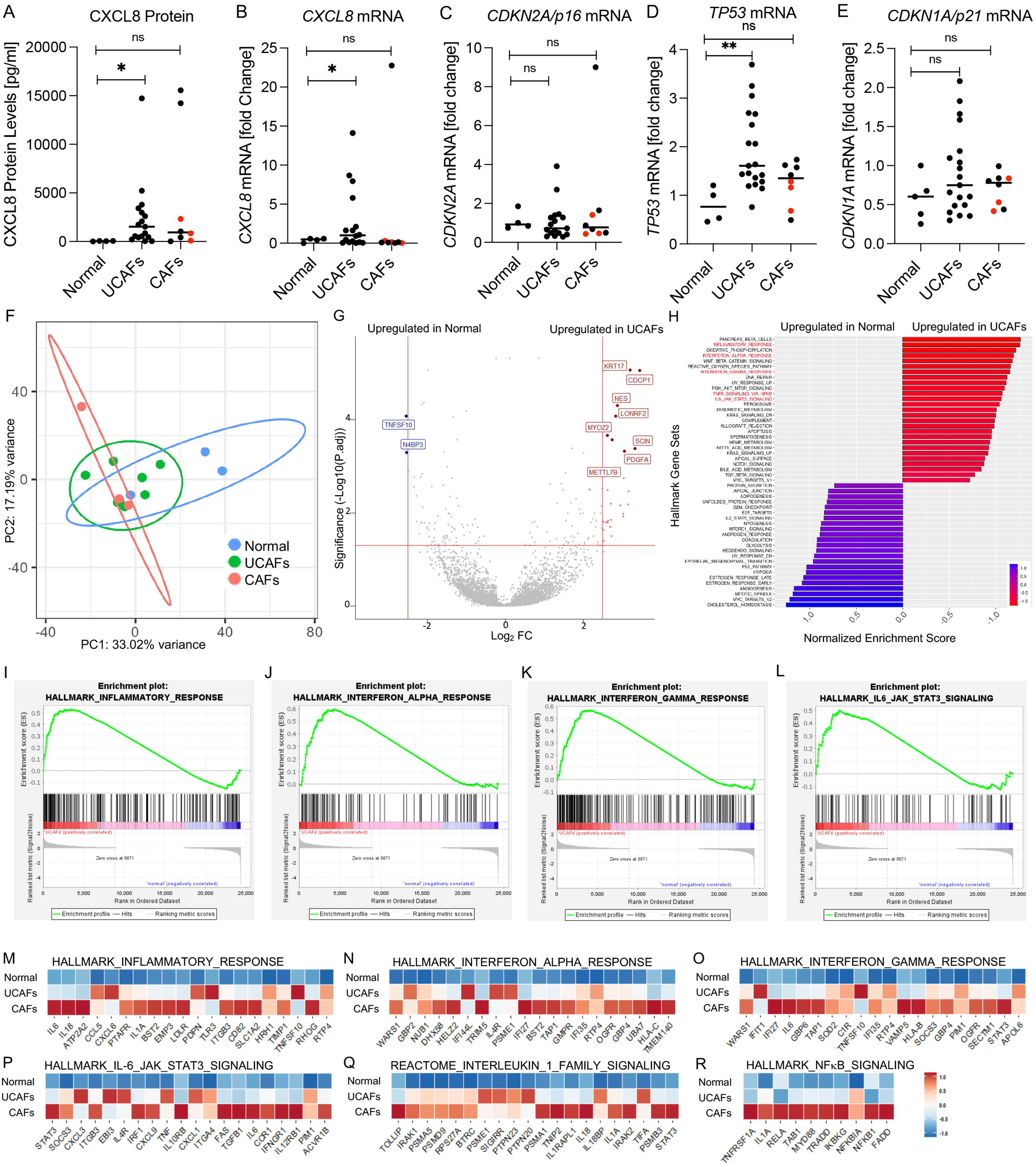
UCAFs and CAFs exhibit an inflammatory, senescence-associated phenotype. (**A**) Enzyme-linked immunosorbents assay (ELISA) analysis of secreted CXCL8 protein present in the conditioned media of human primary colon fibroblasts isolated from the stroma of normal (n = 4), colitis (n = 16), and colon cancer samples (n = 8). Colon cancer samples include colitis-associated cancer (CAC; indicated by the red dots) and sporadic colon cancer samples (indicated by the black dots). (**B-E**) Quantitative real-time PCR (qRT-PCR) relative expression analysis of *CXCL8, CDKN2A, TP53, CDKN1A* mRNAs comparing the same primary fibroblasts as in **(A)**. Each dot represents an individual patient sample. (**F**) Principal component analysis (PCA) of bulk mRNA sequencing comparing normal colon fibroblasts, UCAFs, and CAFs. Ellipsoids show 95% confidence intervals. (**G**) Volcano plot of differentially expressed genes comparing normal *vs.* UCAFs with significance threshold of Log2FC ≥ 2.5 and P.adj < 0.05. (**H**) Gene Set enrichment analysis (GSEA) of the top 25 Hallmark pathways significantly upregulated or downregulated in normal colon fibroblasts compared to UCAFs (**H**). **(I-L)** Enrichment plots for four of the gene sets (indicated in red in **H**). **(M-R)** Heatmaps of the top differentially expressed genes in six of the Hallmark gene sets comparing normal colon fibroblasts (normal), UCAFs, and CAFs. Statistical analysis was performed using Student’s *t*-test. Data are shown as mean ± standard deviation. ns, not significant; *P* >0.05; * *P* ≤ 0.05 and ** *P* ≤ 0.01.

To gain further insight, we then re-analyzed mRNA transcriptome data of normal colon fibroblasts, UCAFs and CAFs that we previously reported.[30] Principal component analysis (PCA) separated UCAFs and CAFs when compared to normal colon fibroblasts with a 33% variance explained by PC1 and a 17.19% variance by PC2 (**Figure 1F**). The volcano plot comparing normal colon fibroblasts and UCAFs (with a 2.5-fold change threshold and an adjusted P value of 0.05) identified 2 genes upregulated in normal colon fibroblasts and 26 in UCAFs (**Figure 1G**). Gene Set Enrichment Analysis (GSEA) using Hallmark gene sets revealed a significant upregulation of inflammatory signaling pathways in UCAFs such as INFLAMMATORY_RESPONSE, INTERFERON _ALPHA_RESPONSE, WNT_BETA_CATENIN_SIGNALING, INTERFERON_GAMMA_RESPONSE, TNFA_SIGNALING_VIA_NFKB, IL6_JAK_STAT3_SIGNALING (**Figure 1H-L** and **Supplementary Figure S1I,J**). In addition, we observed an enrichment in senescence and SASP gene sets (**Supplementary Figure S2A,B**). Similar results were obtained when comparing CAFs to normal colon fibroblasts (**Supplementary Figure S1A-H, Supplementary Figure S2C,D**). Heatmap analyses of the most differentially expressed individual genes within these pathways for all three fibroblast groups demonstrated low expression in normal fibroblasts, intermediate levels in UCAFs, and an further increase in CAFs (**Figure 1M-R**). Together, these data indicate that UCAFs exhibit transcriptional signatures of inflammation and senescence, which are further augmented in CAFs.

### CXCL8 Expression is Regulated by Several Inflammatory Mediators

To determine whether CXCL8 expression is regulated downstream of inflammatory signaling pathways we tested the cytokines (TNFα, IL-1β and IFNγ) as well as the bacterial stimuli, lipopolysaccharide (LPS) and the flagella protein Flagellin (FliC) alone or in combination using the UCAF cell line CT5-S/T and the normal colon fibroblast line CRL1541-S/T. In our analysis, we included the colon cancer epithelial cell line HCT116 as well as HEK293T cells as a non-intestinal control cell type. *CXCL8* mRNA and secreted CXCL8 protein were assessed by qRT-PCR and ELISA, respectively. TNFα induced CXCL8 in all three colon cell types but not the HEK293T cells (**Figure 2A-F** and **Supplementary Figure S3A**). In contrast, IL-1β triggered a strong response in CT5-S/T and the HCT116 epithelial cells, but not the normal colon fibroblasts CRL1541-S/T (**Figure 2A-F** and **Supplementary Figure 3B**). Both fibroblast lines displayed a modest response to LPS treatment alone, whereas HCT116 cells were unresponsive (**Figure 2A-F** and **Supplementary Figure S3C,H**). Conversely, FliC elicited a response only in HCT116 cells, but not in the two fibroblast lines (**Figure 2A-F** and **Supplementary Figure S3D,G**). Combining these treatments (e.g., TNFα and IL-1β) yielded additive and slightly synergistic effects. Finally, IFNγ alone did not induce CXCL8 in any of the cell types, and co-treatment with IFNγ reduced TNFα-induced CXCL8. This effect was not due to a lack in the expression of the respective signaling receptors (**Supplementary Figure S3F**). Instead, it is consistent with the fact that IFNγ signals via the JAK/STAT pathway and not NFκB as the other stimuli tested [33, 34].

**Figure 2:**
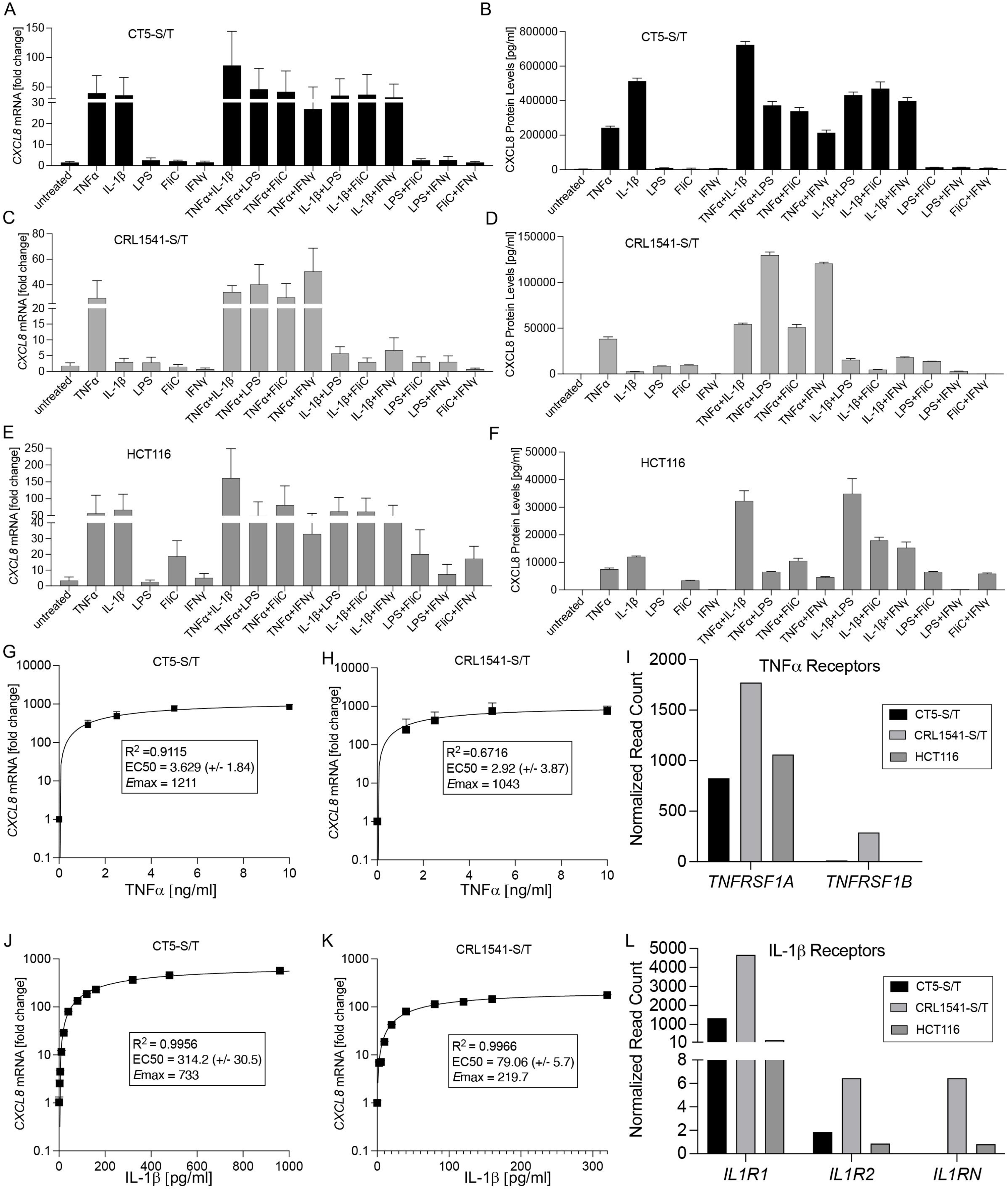
CXCL8 expression is induced by pro-inflammatory cytokines and bacterial signals in colon fibroblasts. **(A-F)** *CXCL8* mRNA and secreted CXCL8 protein expression levels in colitis-associated colon fibroblasts (CT5-S/T), normal colon fibroblasts (CRL1541-S/T) and colon cancer epithelial cells (HCT116) following stimulation with Tumor Necrosis Factor α (TNFα, 20 ng/ml), Interleukin-1 β (IL-1β, 960 pg/ml), Lipopolysaccharide (LPS, 10 μg/ml), Flagellin (FliC,100 ng/ml), and Interferon-γ (IFNγ, 33 ng/ml) either individually or in combination. **(G-H)** Dose-response curves for serial dilutions of TNFα in CT5-S/T and CRL1541-S/T. **(I)** DESeq2-normalized mRNA-seq reads for TNF receptors (*TNFRSF1A* & *TNFRSF1B*) in CT5-S/T, CRL1541-S/T fibroblasts and HCT116 colon cancer epithelial cells. **(J-K)** Dose-response curves for serial dilutions of IL-1β on CT5-S/T and CRL1541-S/T fibroblasts. **(L)** DESeq2-normalized mRNA-seq read counts for the interleukin-1 receptor family members (*IL1R1* & *IL1R2* & *ILRN*) comparing CT5-S/T, CRL1541-S/T fibroblasts and HCT116 colon cancer epithelial cells. Statistical analyses were performed as indicated in each panel. Error bars represent standard deviation of at least three independent experiments. Dose-response curves were obtained by nonlinear regression analysis with R^2^ and EC50 values indicated.

Dose-response assays confirmed that cytokine-induced CXCL8 expression was saturable. TNFα exhibited an EC_50_ of 3.63 ng/ml for the CT5-S/T and 2.92 ng/ml for the CRL1541-S/T (**Figure 2G,H**). This response aligned with the mRNA-seq data of the TNFα receptors (TNFRSF1A and TNFRSF1B), where *TNFRSF1A* was expressed in all three cell types (CT5-S/T, CRL154-S/T and HCT116), while *TNFRSF1B* was expressed predominantly in CRL1541-S/T, but at much lower levels in CT5-S/T and HCT116 (**Figure 2I**). IL-1β EC_50_ values differed between fibroblast lines, with higher sensitivity in the UCAF representative, CT5-S/T (314 pg/ml in) than in normal colon fibroblast CRL1541-S/T (79 pg/ml in) (**Figure 2J,K**). Consistent with this finding, the mRNA-seq data showing differential expression of the IL-1β receptors, IL1R1, IL1R2 & IL1RN (**Figure 2L**). While *IL1R1* and *IL1R2* was expressed in both fibroblast lines (i.e., CT5-S/T and CRL1541-S/T), these receptors were present at lower levels in colon epithelial HCT116 cells. However, the IL-1β receptor antagonist *IL1RN* was primarily expressed in CRL1541-S/T, the normal colon fibroblast, potentially explaining the decreased responsiveness to IL-1β despite higher receptor expression. Together, these data demonstrate that CXCL8 is strongly regulated by inflammatory stimuli in a cell-type and receptor-dependent manner.

### TNFα Accelerates Tumor-Stromal Interactions and Tumor Growth *in vivo*

To address the *in vivo* relevance of the TNFα-mediated inflammatory modulation, we assessed the effect on tumorigenesis in a co-inoculation model (**Figure 3A**). Immunocompromised NSG mice were implanted subcutaneously with a sustained-release pellet of TNFα or a placebo pellet, followed by inoculation of CAC227 epithelial tumor organoids either alone or in combination with the primary UCAF line originally isolated from a colitic colon (CT461-Dist). Tumor growth was monitored by serial caliper measurements.

**Figure 3:**
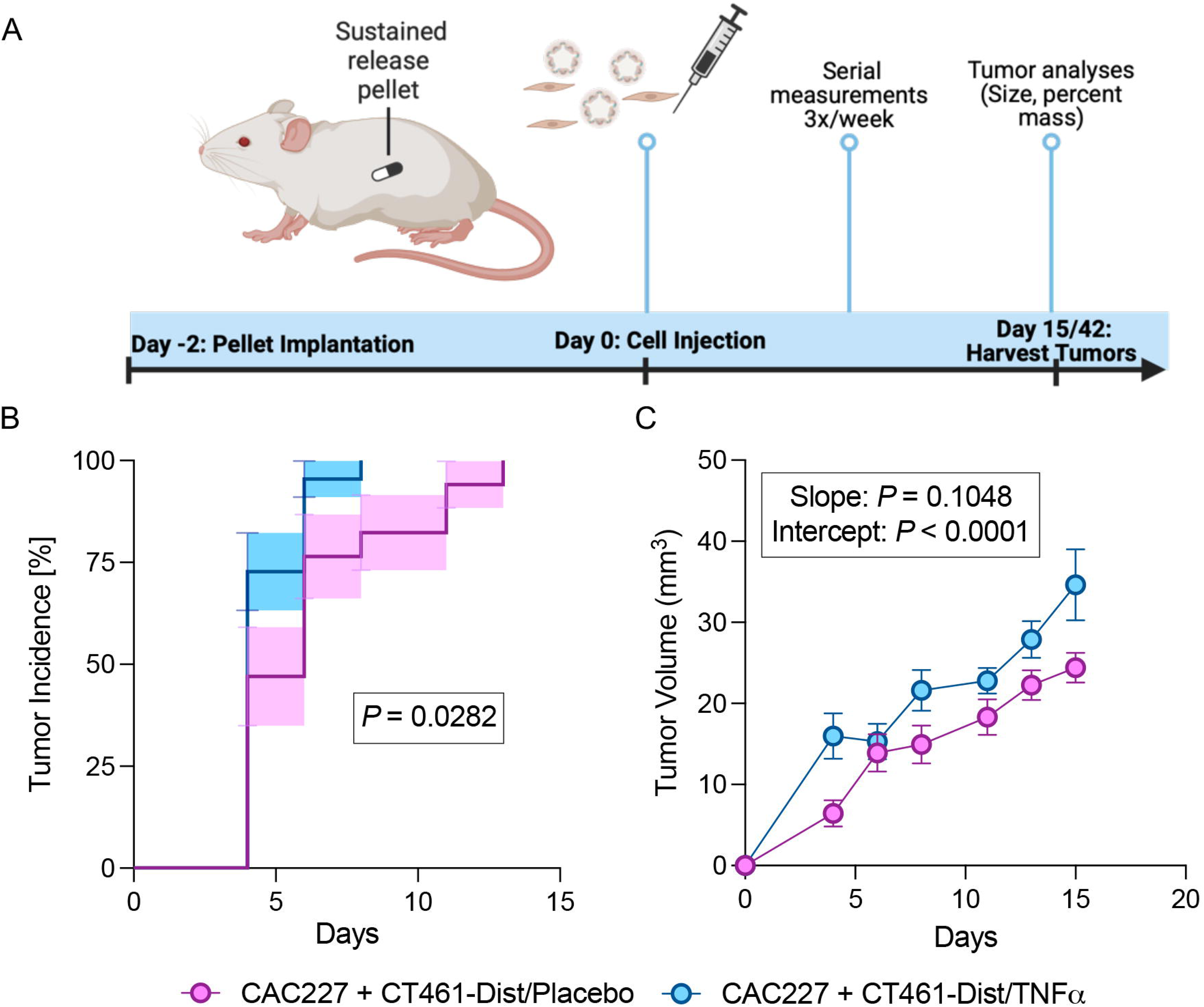
TNFα accelerates tumor growth *in vivo* in a UCAF/CAC co-implantation model. **(A)** Schematic of the *in vivo* study assessing tumor growth by co-injecting UCAFs (CT461-Dist) with CAC epithelial organoids (CAC227) into the flank of immunodeficient non-obese diabetic IL2 gamma-receptor null (NSG) mice. A subset of mice received a slow-release pellet containing TNFα, while control mice received a placebo pellet. Tumor volume was monitored by caliper measurements every 2-3 days and tumors were harvested upon completion for tumor measurements, histology and immunofluorescence analyses. Created with BioRender.com **(B,C**) Tumor latency of the co-inoculated CAC227/CT461-Dist (1×10^4^ cells and 7×10^4^ cells in n=23 and n=22 mice, respectively) is decreased from 6 days in mice implanted with a placebo pellet to 4 days for mice with a slow-release TNFα pellet (*P*=0.0282, Logrank (Mantel-Cox) test, **B**). Similarly, tumor volumes increased from 1.613 mm^3^/day in the mice with the placebo control pellet to 2.013 mm^3^/day in the presence of TNFα pellet (*P*=0.1048 for the slope and *P*<0.0001 for the intercept by linear regression analysis, **C**). Data are shown mean values ± standard error. Statistical analyses were performed as indicated in each panel.

We first assessed whether UCAFs augment tumorigenesis of colon cancer cells in the absence of exogenous TNFα. To this end, we analyzed tumors consisting of CAC227 alone or combined with the CT461-Dist UCAFs in the presence of the placebo pellet. Tumor volume measurements revealed a significant change in the slope (*P* = 0.0319, **Supplementary Figure S4A**) and tumor latency showed a non-significant trend towards reduction from 21 to 14 days (*P* = 0.1701, **Supplementary Figure S4B)**.

Next, we evaluated the impact of TNFα on tumor initiation and progression. In mice co-inoculated with CAC227/CT461-Dist-derived tumors, the addition of a sustained release TNFα-containing pellet reduced tumor latency from 6 days to 4 days when compared to the mice with the placebo pellet (*P* = 0.0282, **Figure 3B**). Tumor growth trajectories were consistent with this reduction in latency. There was no difference in the slope (*P* = 0.10480) but a highly significant difference in the intercept (*P* < 0.0001, **Figure 3C**). Tumor volume increased by 1.61 mm^3^/day in the placebo-treated mice versus, 2.01 mm^3^/day in TNFα-treated mice. These findings were validated in a second independent experiment (**Supplementary Figure S4C**). Finally, at study endpoint, proliferation was assessed by KI67 or PCNA immunostaining, but did not detect differences (**Supplementary Figure S4D, E**) further supporting that TNFα impacts tumor latency, rather than tumor volume (at least at the termination of the study).

Together, these results show that UCAFs enhance tumor initiation, and that TNFα further accelerates this process when UCAFs are present. They highlight a critical role for inflammatory signaling in shaping tumor-stromal interactions *in vivo*.

### Mycoplasma Infections Mimics the Impact of Inflammatory Regulators on CXCL8

The colonic microbiota play a critical role in the pathogenesis of colitis and progression towards CAC [35–38]. Given the technical difficulties in co-culturing bacteria and fibroblasts *ex vivo*, we used the presence of small intracellular bacteria i.e., mycoplasma as a surrogate for microbial crosstalk within the interstitial layer of the colon. As reported previously [39], immunofluorescence and RNAscope^®^ analyses confirmed the presence of mycoplasma in human colon fibroblasts (**Figure 4A** and data not shown).

**Figure 4:**
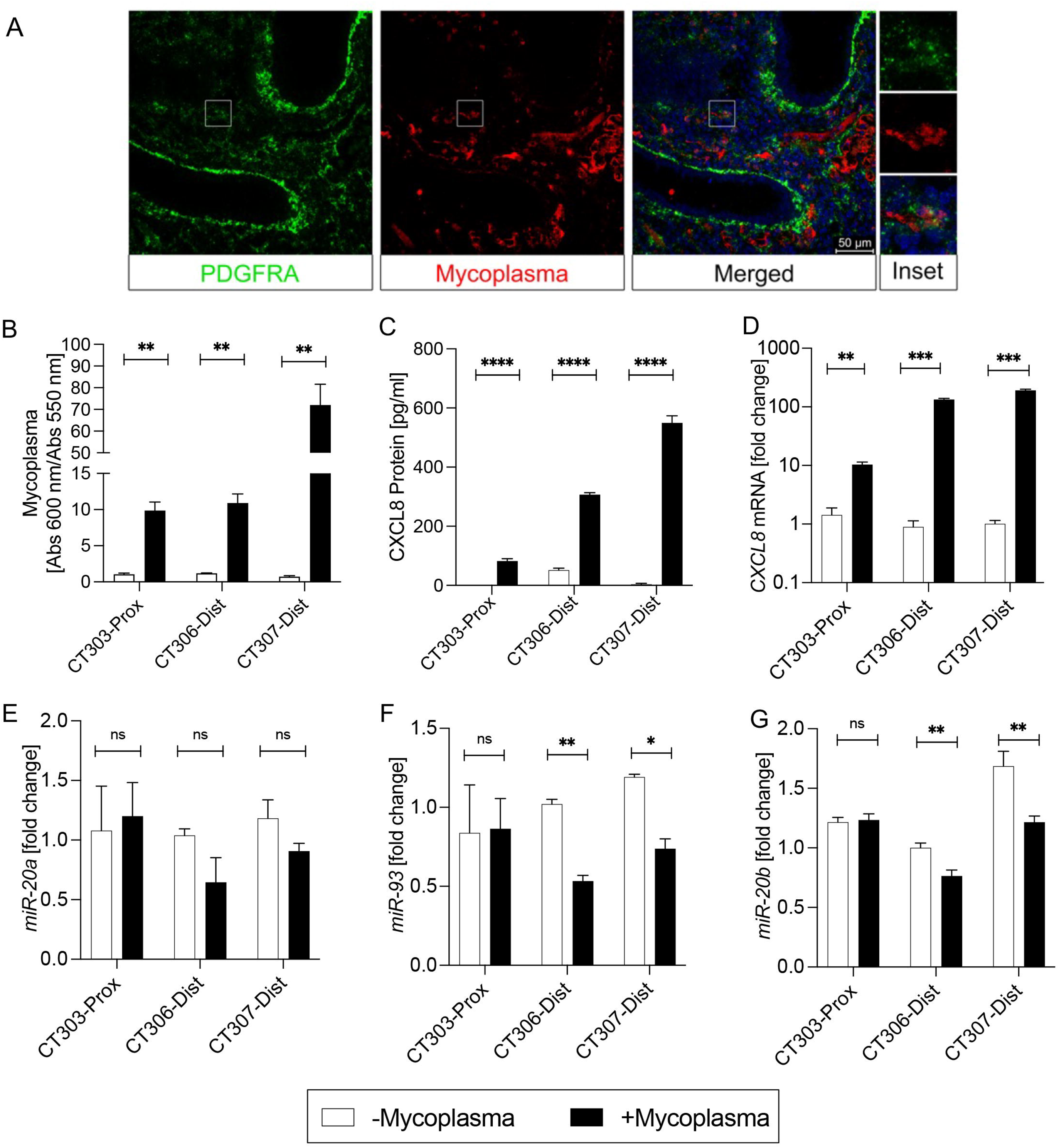
Mycoplasma infection upregulates *CXCL8* expression. **(A)** Immunofluorescence of colon tissue from a patient with ulcerative colitis showing the presence of mycoplasma (red) with the interstitial colon fibroblasts visualized by PDGFRA (green). In the merged panel, nuclei were counterstained with DAPI (blue). (**B)** Mycoplasma colonization status in normal colon fibroblasts (CT303-Prox) and colitis-associated colon fibroblasts (CT306-Dist and CT307-Dist). **(C)** ELISA of secreted CXCL8 protein in matched fibroblasts with or without mycoplasma. **(D-G)** qRT-PCR analysis of *CXCL8* mRNA, *miR-20a, miR-93* and *miR-20b*. Statistical analysis was completed using the student T-test; Data are shown as mean ± standard deviation. ns, not significant, *P* > 0.05; \**P* ≤ 0.01; \*\**P* ≤ 0.001; \*\**P ≤ 0.0001*

During the establishment of the primary fibroblast lines from resected colon tissue, we routinely generate mycoplasma-free cultures. As part of the process, early-passage isolates are cryopreserved prior to decontamination, allowing us to examine paired mycoplasma-positive and mycoplasma-negative cells of the same lines. Using this resource, we compared CXCL8 expression using a normal colon fibroblast line (CT303-Prox) and two UCAF lines (CT306-Dist and CT307-Dist) with and without mycoplasma (**Figure 4B**). All three cell lines exhibited significantly higher levels of CXCL8 protein and mRNA levels in the presence of mycoplasma (**Figure 4C,D**). As already shown in **Figure 1A,B**, baseline CXCL8 levels are elevated in mycoplasma-free UCAFs compared to normal colon fibroblasts. This difference was further enhanced in the presence of mycoplasma (compare CT303-Prox to CT306-Dist and CT307-Dist in **Figure 4C,D**) suggesting a synergistic relationship. In our previous study [30], we discovered that *miR-20a*, a miRNA of the *miR-17* family is a strong regulator of CXCL8 expression. When we examined the expression of the *miR-17* family member, the *miR-20a* and *miR-20b* levels were unchanged when comparing mycoplasma-free normal fibroblasts to UCAFs and CAFs, while *miR-93* was slightly upregulated (**Supplementary Figure S5**). In contrast, all three miRNAs were downregulated in the presence of mycoplasma (**Figure 4E-G**), consistent with their known role in restraining inflammatory transcripts.

To determine whether mycoplasma upregulates CXCL8 expression via TNFα or Il-1β, we assessed the role of NFκB signaling, which is their primary downstream signaling mediator [40]. Indeed, IKK-16, a selective IκB kinase inhibitor, blocked the induction of *CXCL8* mRNA by TNFα in CT5-S/T and CRL1541-S/T cells (**Supplementary Figure S6A**). Treatment of mycoplasma-containing CT303-Prox, CT306-Dist and CT307-Dist with IKK-16, significantly suppressed *CXCL8* mRNA, indicating convergence on NF-κB signaling (**Figure 5A-C**). Furthermore, we performed complementary experiments to address whether the maximum levels of *CXCL8* were altered upon mycoplasma infection. However, adding TNFα or LPS to both mycoplasma-free and mycoplasma-positive colon fibroblasts reached comparable maximal level of *CXCL8* mRNA expression irrespective of mycoplasma levels (**Figure 5D-F**). Together, these experiments support that mycoplasma is an *in vivo* regulator of CXCL8 that primes fibroblasts through NF-κB signaling but does not alter overall signaling capacity.

**Figure 5:**
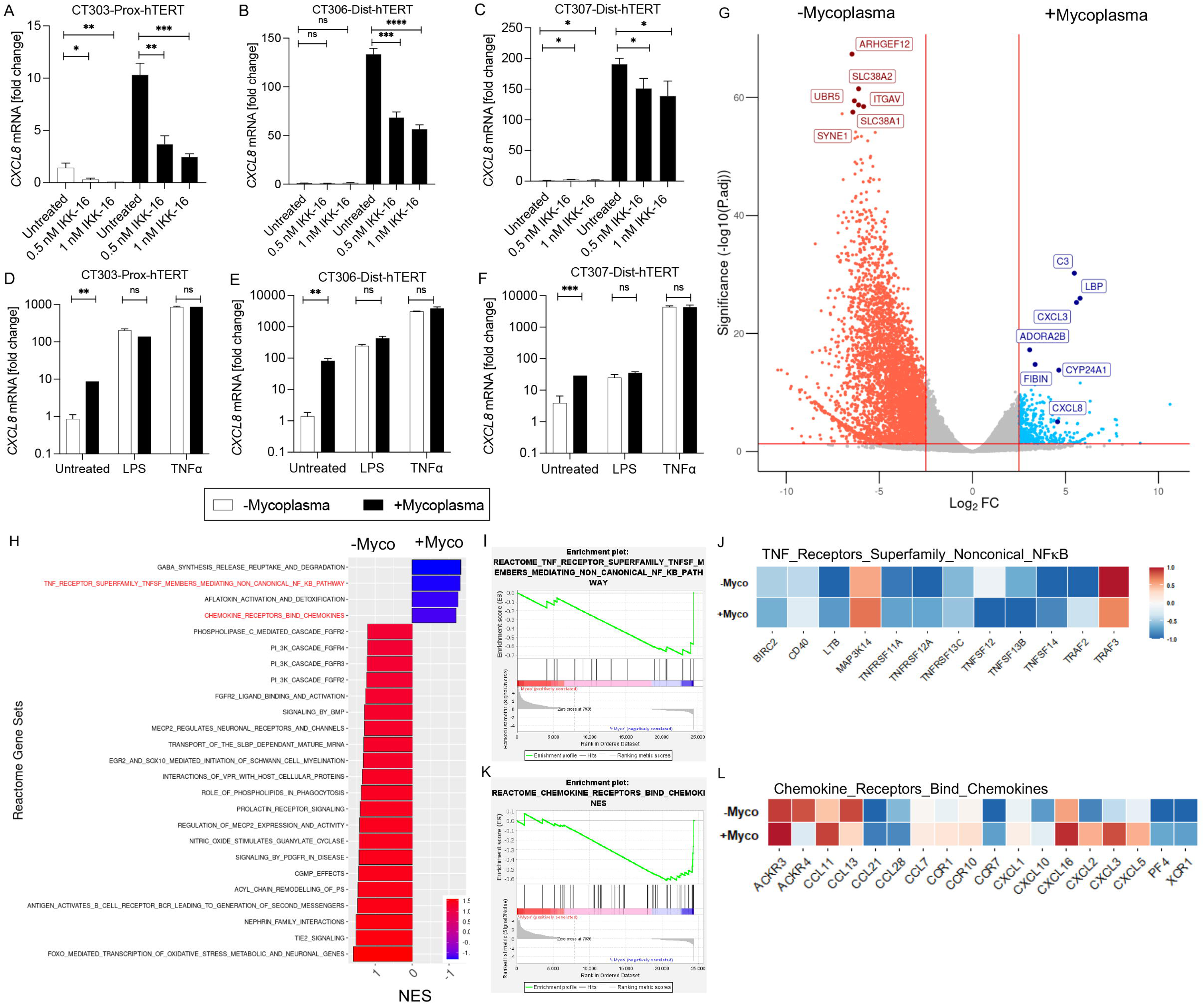
*CXCL8* expression is regulated by NF-κB signaling and modified by mycoplasma colonization. **(A-C)** qRT-PCR of *CXCL8* in a normal colon fibroblast line (CT303-Prox) and two colitis-associated colon fibroblast lines (CT306-Dist & CT307-Dist) in the presence or absence of mycoplasma treated with NF-κB inhibitor, IKK-16 (**D-F**) *CXCL8* mRNA levels following stimulation with TNFα or LPS in the same fibroblast lines. **(G**) Volcano plot comparing gene expression changes as a consequence of mycoplasma with a significance threshold of 2.5 for Log2FC and 0.05 for P.adj. **(H-L)** GSEA depicting the top upregulated and downregulated Reactome gene sets in the mycoplasma-positive and -negative colon fibroblasts (**H**, NES, Normalize Enrichment Score). Enrichment plots and corresponding heatmaps for two gene sets highlighted in red in (H), REACTOME_TNF_RECEPTOR_SUPERFAMILY_TNFSF_MEMBERS_MEDIATING_NON_CANONICAL_NF_KB_PATHWAY **(I-J)** and REACTOME_CHEMOKINE_RECEPTORS_BIND_CHEMOKINES **(K,L)**. All cell culture experiments were performed with at least three biological replicates; data are shown as mean ± standard deviation. Statistical analysis was completed using the Student’s t-test. ns, not significant, *P* > 0.05; * *P* ≤ 0.05; ** *P* ≤ 0.01; *** *P* ≤ 0.001; **** *P* ≤ 0.0001.

Transcriptome analyses further substantiated these results. As expected, mycoplasma infection was the major principal component separating the data (PC1=82.66%, **Supplementary Figure S6C**). In fact, the majority of the observed transcriptional changes was the downregulation of gene expression in mycoplasma-positive fibroblasts (**Figure 5G)**, with only a small subset upregulated including the chemokines *CXCL3* and *CXCL8*. GSEA confirmed widespread downregulation of Hallmark and Reactome gene sets (**Figure 5H** and **Supplementary Figure S6D**). Among the few pathways upregulated were TNF_RECEPTOR_SUPERFAMILY_(TNFSF)_MEMBERS_ MEDIATING_NON-CANONICAL_NF-KB PATHWAY, the CHEMOKINE_RECEPTORS _BIND_CHEMOKINES and the HALLMARK_IL6_JAK_STAT3_SIGNALING (**Figure 5I-L** and **Supplementary Figure S6E,F**) again supporting that inflammatory responses are a major consequence of mycoplasma infection.

Together these data demonstrate that mycoplasma infection can serve as a paradigm for microbial-fibroblast interactions in the inflamed colon. The effect on CXCL8 regulation is mediated through shared NF-κB-dependent pathways, highlighting a microbial mechanism for amplifying stromal inflammatory signaling relevant to colitis.

### Colitis-associated Fibroblasts Retain their CXCL8 Response throughout Stem Cell Reprogramming

We next sought to explore why normal colon fibroblasts and UCAFs respond differently to inflammatory stimuli. We previously reported that the differences between normal, UCAFs and CAFs in respect to CXCL8 secretion are partly mediated by miRNAs of the *miR-17* family [30]. This observation was supported in our current study. Although mycoplasma increased CXCL8 expression in normal colon fibroblasts and UCAFs, *miR-17* family members were consistently downregulated **(Figure 4E-G**). Interestingly, this appears to be independent of the major transcription factor regulating the expression of the *miR-17* family, MYC.[41, 42] qRT-PCR analysis for *MYC* did not show downregulation as a result of the presence of mycoplasma (**Supplementary Figure S6B**).

We also assessed whether inflammatory cytokines and bacterial stimuli could influence expression of miRNAs of the *miR-17* family. Interestingly, expression of *miR-20a* and *miR*-93 was only changed in the CT5-S/T UCAFs, but not normal CRL1541-S/T fibroblasts (**Supplementary Figure S3I-N**). These findings suggested an intrinsic difference between normal colon fibroblasts and UCAFs in the regulation of CXCL8 expression.

Based on these results, we wondered whether this was due to “inflammatory memory”, as previously described in skin fibroblasts.[43] To test this hypothesis we made use of the mesenchymal derived fibroblasts (MDFs) we recently reported [31]. These MDFs are isolated from organoids, which were generated using induced pluripotent stem cells (iPSCs) derived from normal colon fibroblasts or UCAFs (**Figure 6A**). These MDFs preserve the genotype of the original fibroblasts but are reset to a more homogeneous and developmentally naive state and lack any exposure to the tumor microenvironment.

**Figure 6:**
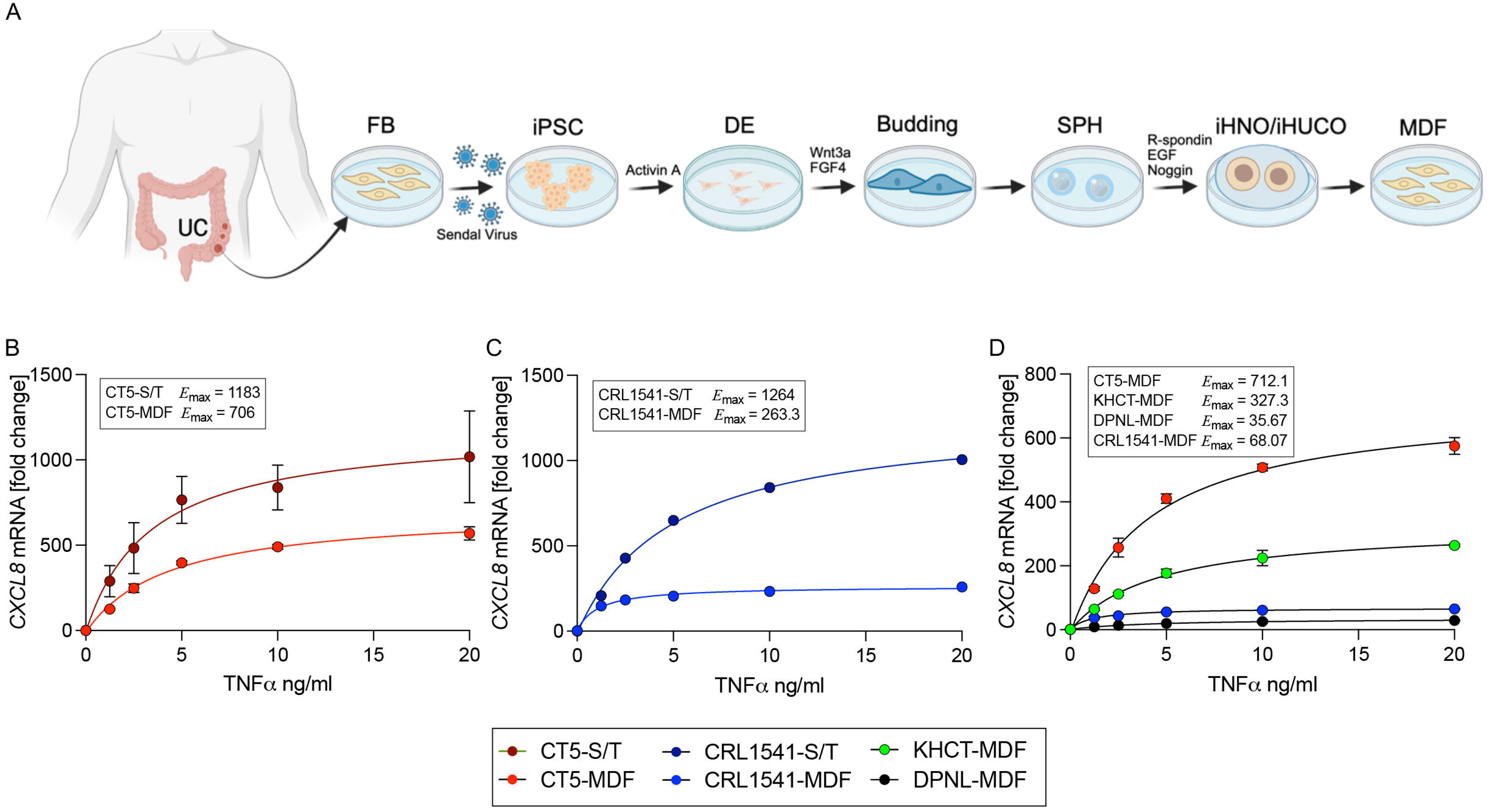
iPSC-derived mesenchymal fibroblasts from colitis retain heightened CXCL8 responsiveness to TNFα. **(A)** Schematic of the generation of Mesenchymal-Derived Fibroblasts (MDFs) from induced pluripotent stem cells (iPSCs) reprogrammed from normal colon fibroblasts and UCAFs. UC, ulcerative colitis; FB, fibroblasts; DE, definitive endoderm; SPH, spheres; iHNO, induced human normal colon organoids; iHUCO, induced human ulcerative colitis organoids; MDF, mesenchymal-derived fibroblasts. **(B,C)** Dose response curves for TNFα stimulation for MDFs derived from CT5 (UCAF) and CRL1541 (normal colon fibroblast) compared to their immortalized parental counterparts. (**D**) Comparative TNFα dose-response of MDFs from iPSC-derived normal colon fibroblasts (CRL1541-MDF, DPNL-MDF) *vs.* UCAFs (CT5-MDF, KHCT-MDF) shows a significantly stronger CXCL8 induction in MDFs derived from UCAFs than normal colon fibroblasts. Data represent the mean of at least three biological replicates; error bars indicate standard deviation. Dose-response curves were obtained by nonlinear regression analysis, with E_max_ values indicated. Differences in E_max_ values were assessed extra sum-of-squares F test; all E_max_ values were significant from each other with *P*<0.0001.

When the CT5-MDFs were compared to their immortalized counterpart, CT5-S/T (**Figure 6B**), CXCL8 induction by TNFα was moderately reduced, with a lower *E*_max_ value (706 *vs.* 1,183) and a slightly higher EC_50_ (4.34 *vs.* 3.39 ng/ml). In contrast, the CRL1541-MDFs displayed a markedly attenuated response compared with CRL1541-S/T fibroblasts (*E*_max_ of 263 *vs.* 1,264, EC_50_ of 1.11 *vs.* 4.92 ng/ml) (**Figure 6C**). Additional MDFs, one from a normal colon fibroblast line (DPNL-MDF) and one from a UCAF line (KHCT-MDF), corroborated the results (**Figure 6D**): MDFs from normal fibroblasts exhibited blunted *CXCL8* induction (*E*_max_ of 35.67 and 68.07), whereas MDFs from UCAFs retained a higher responsiveness (*E*_max_ of 712.1 and 327.3), respectively. Together, these findings demonstrate that - at least in our limited number of samples - iPSC reprogramming of UCAFs retained TNFα responsiveness similar to immortalized UCAFs, while iPSC reprogramming of normal colon fibroblasts resulted in a more attenuated response. This indicates that epigenetic memory or stable transcriptional programming may underlie the persistent inflammatory phenotype observed in the CAFs.

## DISCUSSION

### The Senescence-Associated Secretory Phenotype of Colon Fibroblasts

The focus of this study was to elucidate the mechanisms responsible for the elevated CXCL8 secretion from interstitial fibroblasts that contribute to the progression of ulcerative colitis to CAC, which we previously reported.[23, 30] Using primary colon fibroblast cultures isolated from patients undergoing colon resection and their immortalized derivatives, we conducted a comprehensive analyses across a large cohort to account for biological heterogeneity due to genetic or environmental factors. The primary observation was that the expression profile of UCAFs reflects an intrinsically inflammatory state, based on the upregulation of CXCL8 mRNA and protein expression, and the enrichment of multiple inflammatory signatures, including HALLMARK_ INFLAMATORY_RESPONSES, HALLMARK_INTERFERON_ALPHA_ RESPONSES, and HALLMARK_INTERFERON_GAMMA_RESPONSES.

This pro-inflammatory state is likely triggered by stress-induced senescence as two of the key transcriptional activators in this process *TP53* and *CDKN1A* are upregulated in primary human UCAFs compared to normal colon fibroblasts, consistent with prior descriptions of the SASP.[14] Notably, this senescent phenotype was present in fibroblasts isolated *ex vivo*, suggesting that senescence and inflammation are stable and cell-intrinsic properties of UCAFs. However, this ‘baseline’ state can be further amplified by inflammatory cytokines (e.g., TNFα or IL-1β) or microbial stimuli (e.g., LPS or mycoplasma). Conversely, it can be reduced by inhibition of NFκB signaling, signaling *in vivo*, implantation of a TNFα sustained-release pellet accelerated tumor onset and growth in the presence of UCAFs confirming the pathophysiological relevance of this axis (**Figure 3**). While we have not addressed the precise source of these inflammatory ligands *in vivo*, these likely derive from epithelial, stromal, immune, and microbial interactions within the inflamed colon microenvironment. The cell type-specific responses observed in our study are likely due to the expression levels of the different receptors. While TNFα and IL-1β receptors are expressed in all cell types studied, the LPS receptor TLR4 is only present in the colon fibroblasts used in this study, but not the colon cancer epithelial cell line. As such, its expression parallels the response of these cells to LPS (**Figure 2I,L** and **Supplementary Figure S3H**).

Another interesting observation from the study of the primary colon fibroblasts, is that the progression from a normal colon to colitis and ultimately to CAC is not linear. CAFs do not “just” augment the inflammatory phenotype observed in the UCAFs. While most of the Hallmark gene signatures in the CAFs are even more upregulated compared to the UCAFs (**Figure 1M-R**), individual genes of these gene sets do not all follow this trend. For example, in the HALLMARK_ INFLAMATORY_RESPONSES (**Figure 1M**) multiple genes (*CCF5*, *CXCL6*, *PDPN*, *TLR3* and *TNFSF10*) are high in UCAFs, but reduced in CAFs, suggesting that CAFs may undergo pro-tumorigenic remodeling. Similarly, CXCL8 levels are reduced in CAFs (**Figure 1A,B**). This more nuanced response could be attributed to a change in the microenvironment, where immune cells take over some of the inflammatory signaling originally provided by the fibroblasts. In addition, colon cancer cells release anti-inflammatory factors to remodel the surrounding tissue, to support tumor growth and to evade immune detection. As a result, CAFs must adapt to the changed tumor microenvironment. Indeed, it has been shown that CAFs tend to shift their role to supporting tumor progression and remodeling of the extracellular matrix during disease progression [44].

### Differences Between Normal Colon Fibroblasts and UCAFs

One of the most striking differences between normal colon fibroblasts and UCAFs is the magnitude in CXCL8 induction both at baseline and upon stimulation. This was observed in the collection of the primary colon fibroblasts (**Figure 1A,B**) and upon stimulation with increasing amounts of e.g., IL-1β (**Figure 2J,K**) or by the presence of mycoplasma (**Figure 4C,D**). To understand the difference between normal colon fibroblasts and UCAFs, we first focused on miRNAs. We previously published that miRNAs of the *miR-17* family regulate CXCL8 expression and are downregulated in UCAFs and CAFs when compared to normal colon fibroblasts establishing an inverse relationship [30]. Here we now show that miRNAs of the *miR-17* family are also changed by inflammatory signals or mycoplasma infection. Yet, this appears to be restricted to UCAFs and was not observed in normal colon fibroblasts. Moreover, the extent of the changes in the expression levels of the miRNAs of the *miR-17* family does not seem likely to account for the degree of change in *CXCL8* expression. Direct modulation of the *miR-17* miRNA family expression levels is formally needed to exclude this potential mechanism.

A similar conclusion was obtained for the NFκB signaling pathway, which is one of the main mediators of the SASP in many cell types [32, 45–47]. While the NFκB inhibitor IKK-16 reduced CXCL8 expression, it was not sufficient to completely diminish all expression (**Figure 5A-C**).

Our data argue that other pathways are involved in the regulation of CXCL8 expression. To this end, we considered inflammatory memory, similar to studies shown in other tissues such as skin [43]. We previously reported that iPSC-derived induced human UC-derived organoid from UCAFs retained their functional features compared to an iPSC-derived normal organoid model derived from normal colon fibroblasts [31]. Here we show that iPSC-derived mesenchymal fibroblasts from these UCAF-derived organoids showed a robust CXCL8 response to TNFα, while the iPSC-derived colon fibroblasts from normal colon fibroblasts showed a very limited response. These findings suggest that colon fibroblasts can retain phenotypic features of the original fibroblasts and exhibit a disease memory in how to respond to inflammatory stimuli. While these data are very exciting, additional experiments will be needed to identify the mechanism underlying this memory. Based on the original study in the skin [43], this will likely involve epigenetic changes that are not erased during the establishment of the iPSCs.

### Inflammation as a Ground State for Colon Fibroblasts?

Besides understanding these changes in the fibroblasts upon colitis and their contribution to tumor progression, the study also highlights a technical challenge. The inflammatory status of the actual colon environment in the human body is clearly very different from how cells are maintained in tissue culture. *In vivo*, epithelial cells, fibroblasts, immune cells, microbiota as well as the components of the feces form an active microenvironment, Yet, standard lab cell culture protocols are based on a “clean” environment, where fibroblasts lack any signs of contamination (e.g., mycoplasma) or the presence of bacteria. Our data now suggest that a “clean” UCAF lacks many characteristics of the UCAFs present *in vivo* and is thus less representative of an actual colon environment. Thus, future studies of UCAFs will benefit from experimental systems that better recapitulate the native colonic microenvironment present in the human colon.

## CONCLUSIONS

In summary, our study identified UCAFs and CAFs as key stromal mediators of inflammation in UC and CAC, which are characterized by a stable, cell-intrinsic senescence and persistent CXCL8 expression. We demonstrate that fibroblast-derived CXCL8 expression is amplified by cytokines and microbial signals through NFκB pathways and that this heightened responsiveness is retained even after iPSC reprogramming, suggesting epigenetic memory. These findings highlight fibroblasts not only as contributors to tumor-stroma crosstalk but also as potential biomarkers of disease progression and therapeutic targets for intercepting inflammation-driven colon oncogenesis.

## ACKNOWLEDGMENTS

The authors would like to thank the patients and their families for their contribution to the study. We also would like to thank the members of the Wessely and Huang lab for critical inputs towards the progress of the project. This work was supported by grants from NIH/NCI (7RO1 CA237304) to OW and EH and 5U01CA214300 to EH. HG was supported by a Cleveland Clinic Catalyst grant.

## AUTHOR CONTRIBUTIONS

- Conceptualization and study design: EH, OW
- Acquisition of data/experiments: KA, UT, HG, YB, MB, MA, SKS
- Data analysis and interpretation: KA, UT, RAS, RCF, MCr, MCh, ZC
- Drafting of the manuscript: KA, EH, OW
- Critical revision of the manuscript: KA, UT, EH, OW
- Supervision and funding acquisition: EH, OW
- All authors read and approved the final manuscript

## SUPPLEMENTARY FIGURE LEGENDS

**Supplementary Figure S1: Differential gene expression and enriched pathways in CAFs. (A)**. Volcano plot of differentially expressed genes from mRNA sequencing analysis comparing normal colon fibroblasts and CAFs. **(B)** GSEA comparing the top thirty upregulated Hallmark gene sets in CAFs *vs.* normal colon fibroblasts. (**C-J)** Select GSEA enrichment plots for four of the gene sets indicated in red in (**B**) as well as the Reactome gene set REACTOME_INTERLEUKIN_1_FAMILY_SIGNALING are shown for CAFs *vs.* normal colon fibroblasts (**C-H**) or UCAFs vs. normal colon fibroblasts (**I-J**).

**Supplementary Figure S2: Senescence pathways in normal colon fibroblasts, UCAFs and CAFs. (A-D)** GSEA enrichment plots for SAUL_SEN_MAYO (**A,C**) and REACTOME_SENESCENCE_ASSOCIATED_SECRETORY_PHENOTYPE_SASP (**B,D**) comparing UCAF *vs.* normal colon fibroblasts (**A,B**) and CAFs *vs.* normal colon fibroblasts (**C,D**). **(E,F)** Heatmaps showing the top differentially expressed genes of the two gene sets comparing normal colon fibroblasts (normal), UCAFs, and CAFs.

**Supplementary Figure S3: CXCL8 induction and receptor expression in response to inflammatory stimuli across colon fibroblasts and epithelial cells. (A-E)** qRT-PCR for *CXCL8* mRNA in a UCAF line (CT5-S/T), normal colon fibroblasts (CRL1541-S/T), a colon cancer epithelial cell line (HCT116) and HEK293 following stimulation with serial dilutions of 1.25 - 20 ng/ml TNFα, 80 - 960 pg/ml IL-1β, 0.63 - 10 μg/ml LPS, 6.25 - 100 ng/ml FliC, 2.1 - 33.3 ng/ml IFNγ. **(F-H)** DESeq2-normalized mRNA-seq read counts of IFNγ receptors (*IFNGR1* & *IFNGR2*), the Flagellin receptor, Toll-like receptor 5 (*TLR5*), and the LPS receptor, Toll-like receptor 4 (*TLR4*), in CT5-S/T, CRL1541-S/T fibroblasts and HCT116 colon cancer epithelial cells. (**I-N)** qRT-PCR analysis of *miR-20a*, *miR-93*, and *miR-20b* expression levels in a UCAF line (CT5-S/T) (**I,K,M)** and in a normal colon fibroblast line (CRL1541-ST) (**J,L,N**) treated with 20 ng/ml TNFα, 960 pg/ml IL-1β, 10 μg/ml LPS, 100 ng/ml FliC, 33 ng/ml IFNγ individually or in combination. Data are shown as mean ± standard deviation.

**Supplementary Figure S4: Analysis of tumor growth *in vivo* using a UCAF/CAC co-inoculation model. (A,B)** Co-inoculation of CAC227 epithelial organoids with the CT461-Dist UCAFs (1×10^4^ cells and 7×10^4^ cells, respectively, N=7 mice) compared to CAC227 epithelial organoids alone (1×10^4^ cells, N=5 mice). When compared to the CAC227-only tumors, the tumors formed upon co-inoculation of CAC227 epithelial organoids with CT461-Dist UCAFs exhibited an increase in tumor volume (**A**, *P*=0.0319 using a linear regression analysis), and a trend toward reduced tumor latency (**B**, *P*=0.1701, Logrank (Mantel-Cox) test). Graphs show mean values for each time point and standard error. (**C**) Tumor volumes of the co-inoculated CAC227/CT461-Dist (1×10^4^ cells and 7×10^4^ cells, respectively, N=6 mice each) were increased from 6.219 mm^3^ in the mice with the placebo control pellet to 7.952 mm^3^/day in the presence of TNFα pellet (*P*=0.337 for the slope and P=0.0129 for the intercept by linear regression analysis). Graph shows mean values for each time point and standard error. (**D,E**) Quantification of immunofluorescence for Ki67 (**D**) and PCNA (**E**) comparing the placebo and TNFα slow-release pellet implanted CAC227/CT461-Dist tumors shown in Figure 3B,C when tumors were harvested. Each point is the quantification of an independent section with the standard deviation depicted. Data were compared using Student’s T-test.

**Supplementary Figure S5: Expression of *miR-17* family members in normal colon fibroblasts, UCAFs and CAFs. (A-C)** qRT-PCR of *miR-20a*, *miR-93* and *miR-20b* comparing primary colon fibroblasts isolated from stroma of normal (n = 4), colitis (n = 16), and colon cancer samples (n = 8). The latter contains fibroblasts from CAC (indicated by the red dots) and sporadic colon cancer samples (indicated by the black dots). Each point represents an individual patient-derived fibroblast sample. **(D)** Spearman’s correlation analysis secreted CXCL8 protein *vs. miR-20a* expression levels showed no significant inverse relationship (R^2^= 0.001) Normal colon fibroblasts are indicated by black, UCAFs by red, and CAFs by green dots. Data are shown as mean ± standard deviation. Statistical analysis was performed using Student’s t-test (A-C) and Spearman’s correlation (D). ns, not significant, *P*> 0.05; * *P* ≤ 0.05; ** *P ≤ 0.01*.

**Supplementary Figure S6: NF-κB Signaling Regulates *CXCL8* Expression by Mycoplasma. (A)** qRT-PCR analysis for *CXCL8* mRNA comparing the UCAF line, CT5-S/T, and the normal colon fibroblast line, CRL1541-S/T treated with 0.5 mM or 1 mM IKK-16 in the presence or absence of 20 ng/ml TNFα. **(B)** qRT-PCR analysis of *MYC* mRNA expression in normal colon fibroblasts (CT303-Prox) and UCAFs (CT306-Dist and CT307-Dist) in the presence or absence of mycoplasma. **(C)** Principal component analysis of bulk mRNA sequencing comparing colitis-associated *vs.* normal colon fibroblasts in the presence or absence of mycoplasma. Ellipsoid shows 95% confidence interval. (**D-F)** GSEA of HALLMARK gene sets (**D**) shows that in the presence of mycoplasma only one gene set, HALLMARK_IL6_JACK_STAT3_SIGNALING is significantly upregulated (indicated in red in **D**) with (**E**) showing the GSEA enrichment plot and (**F**) the heatmap of the most differentially expressed genes. Bar graphs show the mean of three biological replicates; data analysis was performed using the student T-test with P>0.05; not-significant, ns.

