## Supplementary Methods for "Inflammatory Stromal Aging in Ulcerative Colitis and Colitis-Associated Cancer"

### Mouse Tumor Experiments

Sustained release pellets lasting for 21-days containing TNFα (#300-01A, Peprotech; 0. 21 μg/pellet) and placebo pellets (Innovative Research of America, Sarasota, Florida) were surgically implanted into the subcutaneous flank in NSG mice and secured in place with a single absorbable 4-0 Vicryl suture. Two days following insertion, cellular co-cultures, consisting of $8x10^3$-$10^4$ epithelial organoids using the CAC isolate CAC227 with $7x10^4$ the primary CT461-Dist (UCAFs originally isolated from a colitic colon) were injected within 0.5 cm adjacent to the pellets. Measurements using calipers then proceeded every 2-3 days. To determine time and treatment effect on tumor volume, multiple linear regression models with interaction were fitted and the models were selected using adjusted $R^2$ and Akaike Information Criterion (AIC) values. All analyses were conducted using R (version 4.4.1) and GraphPad Prism (version 10). Statistical significance was defined as $P < 0.05$.

### TNFa, IL-1β, IFNγ, LPS and FliC treatment

Colon fibroblasts were treated with 20 ng/ml TNFα (#300-01A, Peprotech), 960 pg/ml IL-1β (#200-01b, Peprotech), 33 ng/ml IFNγ (#300-02, Peprotech), 10 ng/ml LPS solution (#00-4976, Invitrogen), and 100 ng/ml FliC (#LS-G3927, LSBio) in DMEM supplemented in 10% FBS for 24 hours, unless otherwise indicated. Cells were re-treated after 24 hours for a total exposure period of 48 hours. Samples were compared to untreated controls.

### CXCL8 ELISA

To determine the amount of CXCL8 in the fibroblast conditioned media, $10^5$ fibroblasts were cultured in 1 ml of DMEM without FBS for 24 hours. The conditioned media was collected and

CXCL8 concentration was determined using the CXCL8 ELISA kit (#ELH-IL8, Raybiotech) according to the manufacturer's instructions.

**RNA isolation, cDNA preparation, qRT-PCR and mRNA sequencing**

Total RNA was extracted using the TRIzol RNA isolation reagent (Invitrogen #15596026) according to manufacturer's protocol. Purity and integrity of the isolated RNA was evaluated by NanoDrop 2000. cDNA was prepared using M-MLV Reverse Transcriptase (#M170A, Promega), random hexamer primer (#SO142, Thermo Fisher Scientific), dNTPs (#R0181, Thermo Fisher Scientific) and Protector RNase Inhibitor (#03335402001, Roche). TaqMan Fast Advanced Master Mix (#4444557, Thermo Fisher Scientific) was used to perform qRT-PCR. Real time primers were obtained from Thermo Fisher Scientific, CXCL8 FAM (#Hs00174103_m1), CDKN1A/p21 FAM (#Hs00355782_m1), CDKN2A/p16 FAM (#Hs00923894_m1), TP53 FAM (#Hs01034249_m1), MYC FAM (#Hs00153408_m1) and GAPDH VIC (#4326317E).

For microRNA (miRNA) quantification the RNA was processed using the TaqMan advanced miRNA cDNA synthesis kit (#A28007, Thermo Fisher Scientific) for poly(A) tailing, adaptor ligation, and reverse transcription. PCR was performed following the manufacture's protocol using the TaqMan Fast Advanced Master Mix (#4444557, Thermo Fisher Scientific) and the following primer sets: *hsa-miR-20a-5p* (#478586_mir*), hsa-miR-20b-5p (#477804_mir), hsa-miR-93-5p (#478210_mir)* as well as *hsa-miR-191-5p (#477952_mir)* as the endogenous control.

For the transcriptomic analysis of the mycoplasma-containing colon fibroblasts library generation and bulk next-generation mRNA sequencing was performed using TruSeq RNA Library Prep Kits (Illumina, San Diego, CA, USA) and the Illumina platform by Psomagen, Inc. (Rockville, MD, USA).

Transcript were aligned using STAR and then transcript abundances were estimated using Salmon [1] against an index of coding sequences from the Ensembl GRCh38 assembly. Transcript-level abundance was imported and count and offset matrices generated using the tximport R/Bioconductor package [2]. Differential expression analysis was performed using the DESeq2 (v1.44.0) [3]. Principal Component Analysis (PCA) was performed using pcaExplorer (v2.30.0), considering the top 500 genes [4]. GSEA was performed using the Molecular Signature Database (MSigDB) as described [5]. GSEA bar graphs were created in R (v4.4.1) using GGplot2 (v3.5.2), and Tidyverse (v2.0.0). The transcriptomic data were submitted to the NCBI's Gene Expression Omnibus (GEO) sequence repository and are available under GEO Series accession number GSE307278. The transcriptomes from the normal primary human colon fibroblasts, UCAFs and CAFs were reported previously [6] and are available under GSE106119.

**Mycoplasma**

To determine the presence of mycoplasma in the fibroblasts culture supernatants MycoAlert Plus mycoplasma detection assays kit (#LT07-703, Lonza) and MycoAlert assay control set (#LT07-518, Lonza) was used following the manufacturer's instructions. A ratio of Read B/Read A > 1.2 indicated substantial mycoplasma contamination. To eliminate mycoplasma, a combination of two antibiotics (BM-Cyclin, #10799050001, Roche) according to the manufacturer's protocol was used for 21 days, fibroblasts were re-tested, and the process repeated until no mycoplasma was detected. Detection of mycoplasma in paraffin-embedded sections was performed by immunofluorescence using the polyclonal anti-mycoplasma pneumoniae antibody (#PA1-7232, Thermo-Fisher), the fibroblast-specific rabbit monoclonal anti-PDGFRA (#3174, Cell Signaling Technology) and Alexa-Fluor™ secondary antibodies.

**Organoid generation and propagation**

Epithelial organoids were generated as described in Sarvestani *et al.*[7] and propagated in Matrigel beads in L-WRN conditioned media.[8] For the co-culture experiments the organoids were released from the Matrigel using Cell Recovery solution, trypsinized, and strained for single cell solutions using Flowmi® 40 µm filters (#BAH136800040, Sigma).

**Mesenchyme Derived Fibroblasts (MDF) Generation**

The organoid-derived mesenchymal cells were established from normal and UC tissue as previously described.[7] Briefly, colon fibroblasts were reprogrammed to iPSCs using UCAFs isolated from 6 patients with established chronic colitis, 4 from patients with normal colonic fibroblasts and one from a commercially available source. Pluripotency of the generated iPSCs was confirmed. Next, we applied an established protocol for intestinal organoid generation[9] to direct differentiation of the iPSCs into definitive endoderm (DE), followed by intestinal spheroid formation. Normal and UC spheroids were cultured in Matrigel for 21 days to develop induced human normal organoids (iHNOs) and induced human ulcerative colitis organoids (iHUCOs), respectively. Once established, to obtain the MDFs, organoids were trypsinized, triturated and placed into culture on tissue-culture treated culture dishes in DMEM (Gibco) supplemented with 10% FBS (Atlanta Biologics) and 1% penicillin/streptomycin (Gibco).
