## Supplementary Figures for "Inflammatory Stromal Aging in Ulcerative Colitis and Colitis-Associated Cancer"

Figure S1 Almotah, et al.

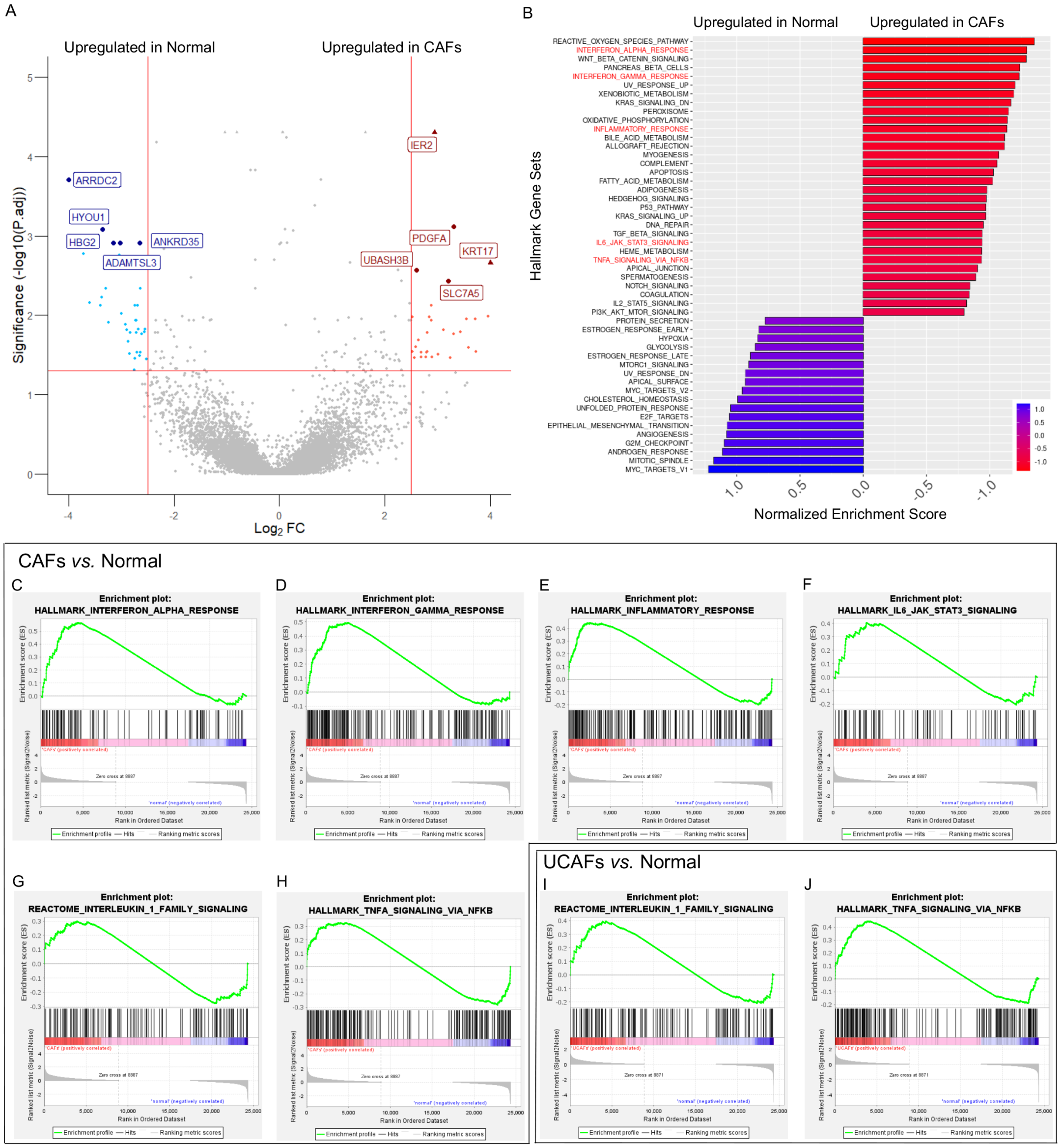


**Supplementary Figure S1: Differential gene expression and enriched pathways in CAFs. (A)**. Volcano plot of differentially expressed genes from mRNA sequencing analysis comparing normal colon fibroblasts and CAFs. **(B)** GSEA comparing the top thirty upregulated Hallmark gene sets in CAFs *vs.* normal colon fibroblasts. (**C-J)** Select GSEA enrichment plots for four of the gene sets indicated in red in (**B**) as well as the Reactome gene set REACTOME_INTERLEUKIN_1_FAMILY_SIGNALING are shown for CAFs *vs.* normal colon fibroblasts (**C-H**) or UCAFs vs. normal colon fibroblasts (**I-J**).

Figure S2   Almotah, et al.

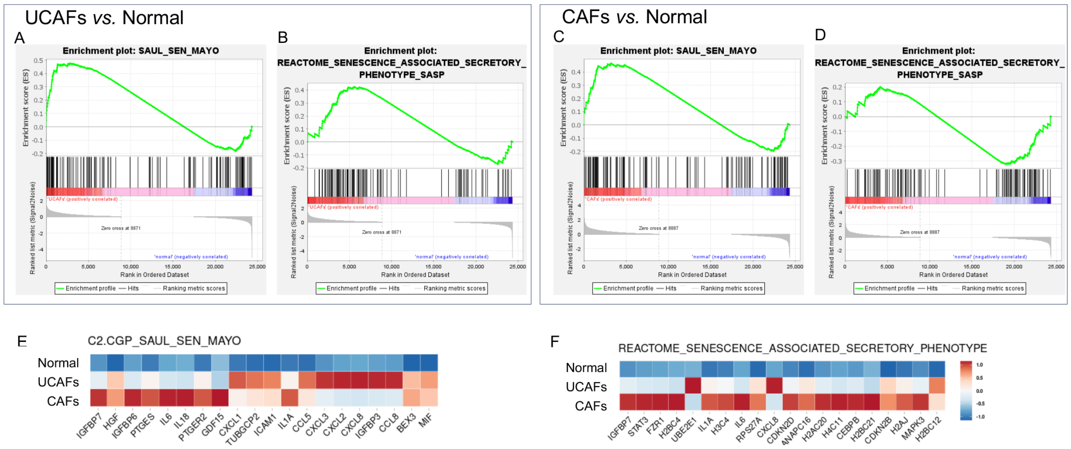


**Supplementary Figure S2: Senescence pathways in normal colon fibroblasts, UCAFs and CAFs. (A-D)** GSEA enrichment plots for SAUL_SEN_MAYO (**A,C**) and REACTOME_SENESCENCE_ASSOCIATED_SECRETORY_PHENOTYPE_SASP (**B,D**) comparing UCAF *vs.* normal colon fibroblasts (**A,B**) and CAFs *vs.* normal colon fibroblasts (**C,D**). **(E,F)** Heatmaps showing the top differentially expressed genes of the two gene sets comparing normal colon fibroblasts (normal), UCAFs, and CAFs.

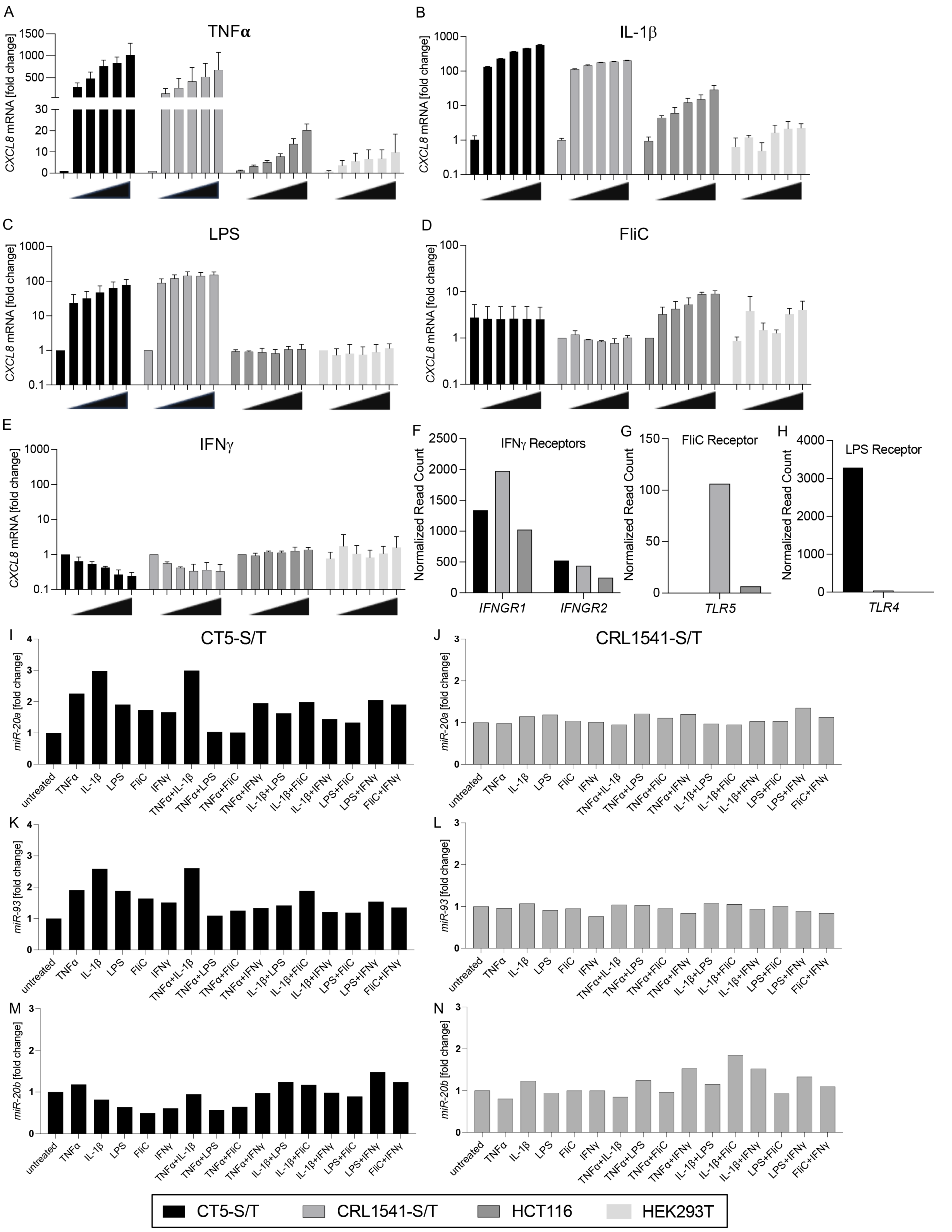


**Supplementary Figure S3: CXCL8 induction and receptor expression in response to inflammatory stimuli across colon fibroblasts and epithelial cells. (A-E)** qRT-PCR for *CXCL8* mRNA in a UCAF line (CT5-S/T), normal colon fibroblasts (CRL1541-S/T), a colon cancer epithelial cell line (HCT116) and HEK293 following stimulation with serial dilutions of 1.25 - 20 ng/ml TNFα, 80 - 960 pg/ml IL-1β, 0.63 - 10 μg/ml LPS, 6.25 - 100 ng/ml FliC, 2.1 - 33.3 ng/ml IFNγ. **(F-H)** DESeq2-normalized mRNA-seq read counts of IFNγ receptors (*IFNGR1* & *IFNGR2*), the Flagellin receptor, Toll-like receptor 5 (*TLR5*), and the LPS receptor, Toll-like receptor 4 (*TLR4*), in CT5-S/T, CRL1541-S/T fibroblasts and HCT116 colon cancer epithelial cells. (**I-N)** qRT-PCR analysis of *miR-20a*, *miR-93*, and *miR-20b* expression levels in a UCAF line (CT5-S/T) (**I,K,M)** and in a normal colon fibroblast line (CRL1541-ST) (**J,L,N**) treated with 20 ng/ml TNFα, 960 pg/ml IL-1β, 10 μg/ml LPS, 100 ng/ml FliC, 33 ng/ml IFNγ individually or in combination. Data are shown as mean ± standard deviation.

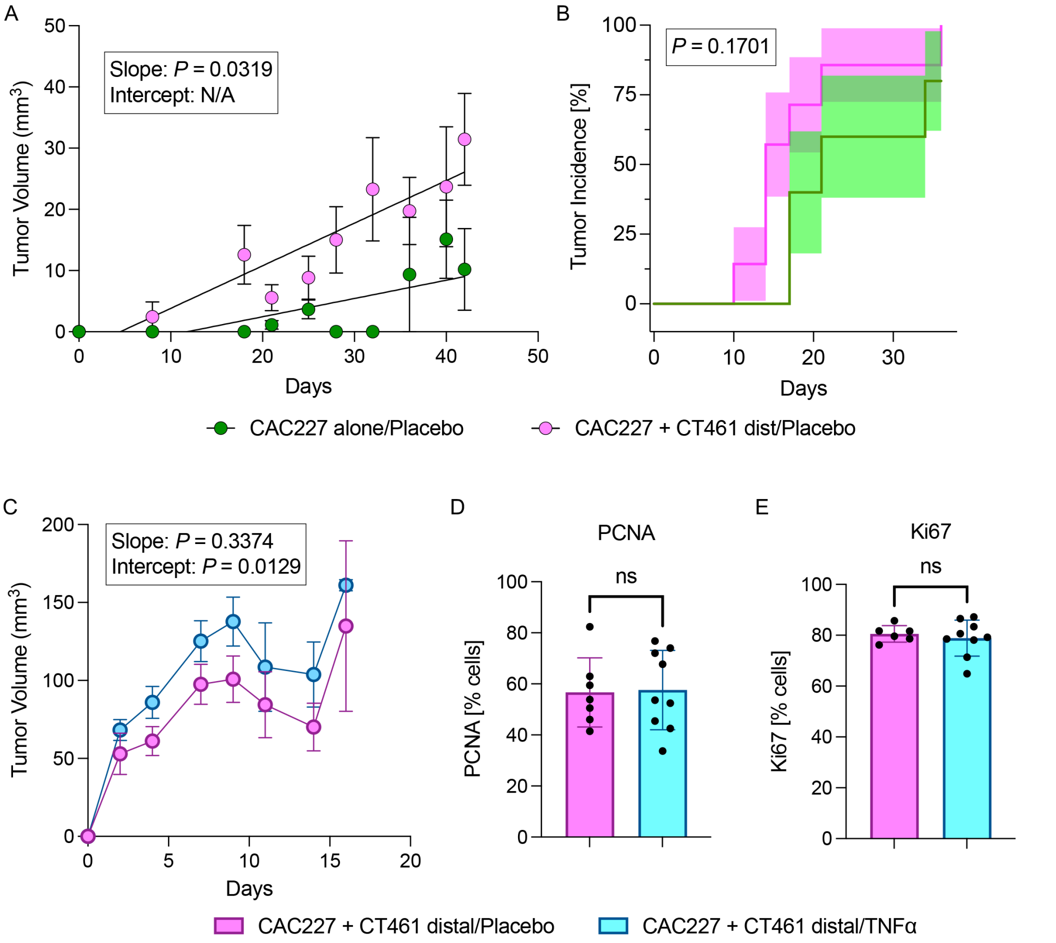


**Supplementary Figure S4: Analysis of tumor growth *in vivo* using a UCAF/CAC co-inoculation model.** **(A,B)** Co-inoculation of CAC227 epithelial organoids with the CT461-Dist UCAFs ($1x10^4$ cells and $7x10^4$ cells, respectively, N=7 mice) compared to CAC227 epithelial organoids alone ($1x10^4$ cells, N=5 mice). When compared to the CAC227-only tumors, the tumors formed upon co-inoculation of CAC227 epithelial organoids with CT461-Dist UCAFs exhibited an increase in tumor volume (**A**, *P*=0.0319 using a linear regression analysis), and a trend toward reduced tumor latency (**B**, *P*=0.1701, Logrank (Mantel-Cox) test). Graphs show mean values for each time point and standard error. (**C**) Tumor volumes of the co-inoculated CAC227/CT461-Dist ($1x10^4$ cells and $7x10^4$ cells, respectively, N=6 mice each) were increased from 6.219 $mm^3$ in the mice with the placebo control pellet to 7.952 $mm^3$/day in the presence of TNFα pellet (*P*=0.337 for the slope and P=0.0129 for the intercept by linear regression analysis). Graph shows mean values for each time point and standard error. (**D,E**) Quantification of immunofluorescence for Ki67 (**D**) and PCNA (**E**) comparing the placebo and TNFα slow-release pellet implanted CAC227/CT461-Dist tumors shown in Figure 3B,C when tumors were harvested. Each point is the quantification of an independent section with the standard deviation depicted. Data were compared using Student's T-test.

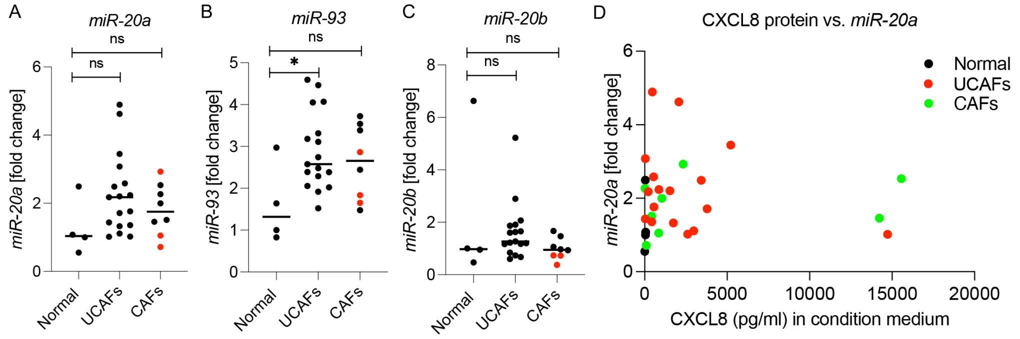


**Supplementary Figure S5: Expression of *miR-17* family members in normal colon fibroblasts, UCAFs and CAFs. (A-C)** qRT-PCR of *miR-20a*, *miR-93* and *miR-20b* comparing primary colon fibroblasts isolated from stroma of normal (n = 4), colitis (n = 16), and colon cancer samples (n = 8). The latter contains fibroblasts from CAC (indicated by the red dots) and sporadic colon cancer samples (indicated by the black dots). Each point represents an individual patient-derived fibroblast sample. **(D)** Spearman's correlation analysis secreted CXCL8 protein *vs. miR-20a* expression levels showed no significant inverse relationship ($R^2$= 0.001) Normal colon fibroblasts are indicated by black, UCAFs by red, and CAFs by green dots. Data are shown as mean ± standard deviation. Statistical analysis was performed using Student's t-test (A-C) and Spearman's correlation (D). ns, not significant, $P$> 0.05; * $P \leq 0.05$; ** $P \leq 0.01$.

Figure S6   Almotah, et al.

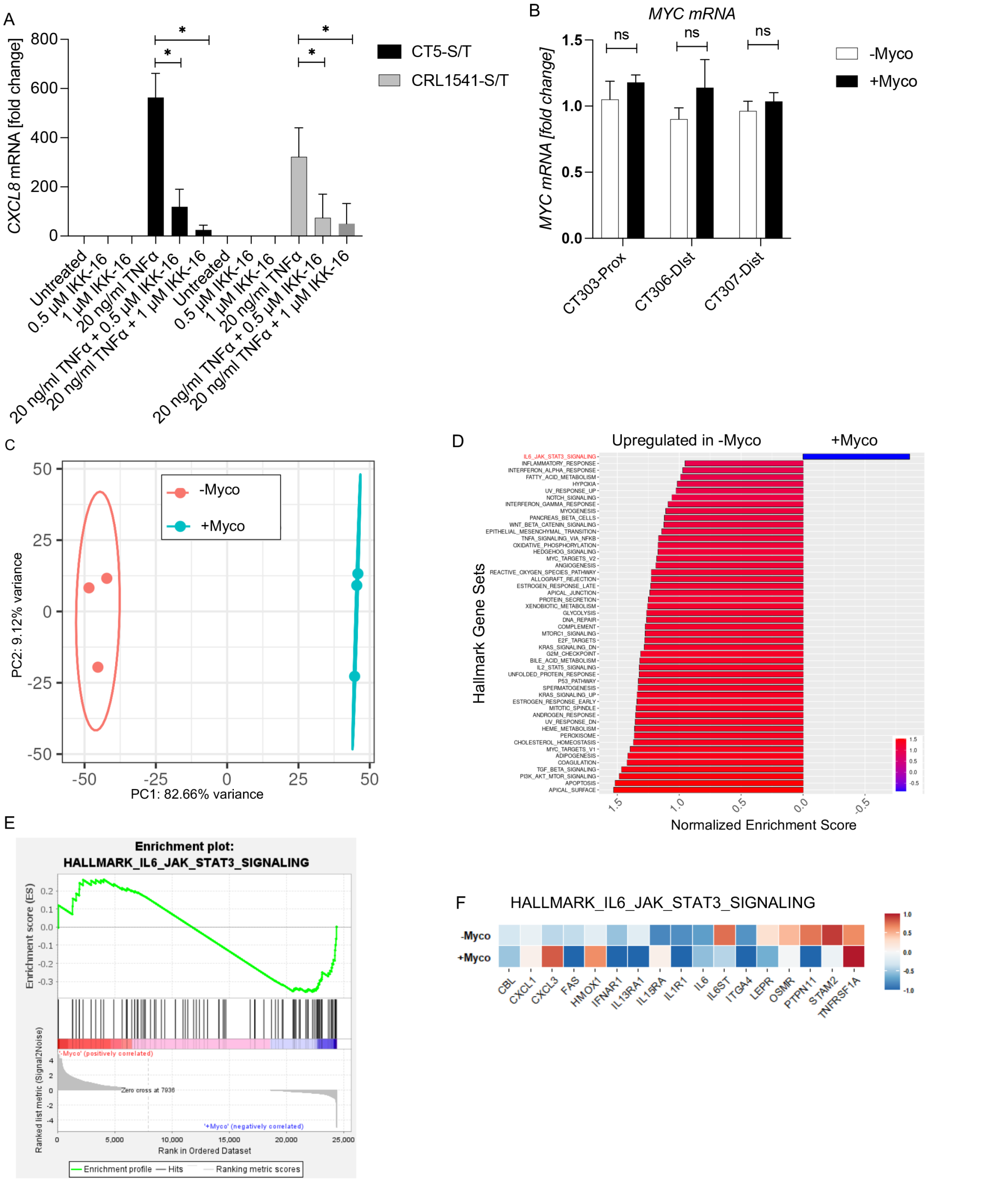


**Supplementary Figure S6: NF-κB Signaling Regulates *CXCL8* Expression by Mycoplasma. (A)** qRT-PCR analysis for *CXCL8* mRNA comparing the UCAF line, CT5-S/T, and the normal colon fibroblast line, CRL1541-S/T treated with 0.5 mM or 1 mM IKK-16 in the presence or absence of 20 ng/ml TNFα. **(B)** qRT-PCR analysis of *MYC* mRNA expression in normal colon fibroblasts (CT303-Prox) and UCAFs (CT306-Dist and CT307-Dist) in the presence or absence of mycoplasma. **(C)** Principal component analysis of bulk mRNA sequencing comparing colitis-associated *vs.* normal colon fibroblasts in the presence or absence of mycoplasma. Ellipsoid shows 95% confidence interval. (**D-F)** GSEA of HALLMARK gene sets (**D**) shows that in the presence of mycoplasma only one gene set, HALLMARK_IL6_JACK_STAT3_SIGNALING is significantly upregulated (indicated in red in **D**) with (**E**) showing the GSEA enrichment plot and (**F**) the heatmap of the most differentially expressed genes. Bar graphs show the mean of three biological replicates; data analysis was performed using the student T-test with P>0.05; not-significant, ns.
