## Supplementary Tables for "Inflammatory Stromal Aging in Ulcerative Colitis and Colitis-Associated Cancer"

| Cell Name | Age (at resection) | Gender | Race | Pathology |
|---|---|---|---|---|
| CAC227-distal | 65 | female | White | Stage 2 cancer; Colitis associated cancer; Colitis |
| CAC227-proximal | 65 | female | White | Colitis |
| CAC231 | 35 | female | white | Stage 3 cancer, colitis associated cancer |
| CAC282 | 65 | Male | White | Chronic and focal active colitis; T1 NO cancer in sigmoid |
| SPCA | 67 | Female | White | Stage 1 colon cancer |
| WGS-CA (CA36) | 62 | Female | white | Stage 1 colon cancer |
| JP-CA | 53 | Male | White | Stage 2 cancer |
| JW-CA | 51 | Male | White | May be rectal cancer - vials identified no STR or pathology (from 2006, likely a match to a cancer) |
| CRL7213 | 46 | Female | White | Colorectal Cancer |
| AACT (CT42) | 49 | Female | White | Diffuse Chronic Active Colitis |
| JW-NL | 51 | Male | White | Ask RF where he found this |
| TVNL | 44 | Male | White | Ask RF where he found this (from sometime between 2007 and 2014, no STR or pathology identified, likely match to a cancer) |
| NL322 | 49 | Male | African American | normal |
| PKNL (NL33) | 62 | Female | White | normal |
| CT5-SV40 | 20 | Male | White | active UC |
| CRL1541-SV40 | 22 weeks | Female | Unknown | normal |
| CT225-distal | 40 | Male | White | Ulcerative colitis |
| CT225-proximal | 40 | Male | White | Ulcerative colitis |
| CT227-distal | 65 | Female | White | First: Normal. Last: Normal. |
| CT227-proximal | 65 | Female | White | First: Normal. No surface epithelium. Last: Normal. No surface epithelium. |
| CT279 distal | 39 | Male | White | Pathologic changes consistent with active Crohn's disease with low-grade dysplasia |
| CT280-distal | 31 | Female | White | Ulcerative colitis |
| CT281-distal | 43 | Male | White | quiescent idiopathic UC; focal low-grade dysplasia |
| CT283-distal | 67 | Male | White | Crohn's Colitis; chronic; patchy active colitis |
| CT284-distal | 58 | Male | White | low grade dysplasia |
| CT284-proximal | 58 | Male | White | high-grade dysplasia in specimen |
| CT285-distal | 57 | Male | White | low grade dysplasia |
| CT285-proximal | 57 | Male | White | ulcerative colitis with focus of low-grade dysplasia |
| CT303-distal | 44 | Male | White | Paraffin: normal; OCT first: normal. OCT last: chronic active colitis, moderate |
| CT303-proximal | 44 | Male | White | normal |
| CT333-distal | 24 | Male | White | Prox: chronic active colitis, severe, Distal: chronic active colitis, moderate |
| CT333-proximal | 24 | Male | White | |
| CT319-Prox | 49 | Male | White | chronic active colitis, severe |
| CT319-distal | 49 | Male | White | Paraffin: normal; OCT first: chronic active colitis, moderate. OCT last: chronic active colitis, moderate. |
| CT320-distal | 39 | Female | White | Paraffin: chronic inactive colitis; OCT last: normal. |
| CT320-proximal | 39 | Female | White | Paraffin: chronic inactive colitis; OCT last: normal. |
| CT306-Proximal | 76 | Female | White | normal |
| CT306-distal-Tert | 76 | Female | White | No active colitis: very little epithelium |
| CT307-distal-Tert | 36 | Male | White | Paraffin: chronic active colitis, severe; OCT first: chronic active colitis, moderate. OCT last: chronic active colitis, moderate |
| CT307-proximal-Tert | 36 | Male | White | normal |
| CT350 prox | 24 | Male | White | Distal: chronic active colitis, moderate, Prox: chronic active colitis, severe |
| CT350 dist | 24 | Male | White | |
| CT351 Prox | 23 | Male | White | Prox and Dist: chronic active colitis, severe |
| CT351 dist | 23 | Male | White | |
| HCT-116 (CCL-247) | Unknown | Male | Unknown | Colorectal Cancer |
| CT420C Prox | 62 | Male | White | Chronic active colitis, moderate |
| CT420C Distal | 62 | Male | White | Chronic active colitis, severe |
| CA404C | 50 | Female | Declined | Invasive adenocarcinoma, moderately differentiated |
| CT461-Proximal | 41 | Female | Indian (Asian) | Prox: chronic active colitis, mild CAC: invasive adenocarcinoma, poorly differentiated with mucinous features |
| CT461-Distal | 41 | Female | Indian (Asian) | Distal: Normal with epi |

Supplementary Table S1: Demographics

| Cell Name | D13S317 | D16S539 | CSF1PO | THO1 | vWA | D21S11 | D7S820 | D5S818 | TPOX | AMEL | D1S1656 | D2S441 | D2S1338 | D3S1358 | D8S1179 | D10S1248 | D12S391 | D18S51 | D19S433 | D22S1045 | DYS391 | FGA | PentaD | PentaE |
|---|---|---|---|---|---|---|---|---|---|---|---|---|---|---|---|---|---|---|---|---|---|---|---|---|
| CAC227-distal | 11, 12 | 12 | 11, 12 | 8, 9.3 | 16, 17 | 28, 30 | 10, 12 | 12 | 8 | X, Y | N.D. | N.D. | N.D. | N.D. | N.D. | N.D. | N.D. | N.D. | N.D. | N.D. | N.D. | N.D. | N.D. | N.D. |
| CAC227-proximal | 11, 12 | 12 | 11, 12 | 8, 9.3 | 16, 17 | 28, 30 | 10, 12 | 12 | 8 | X, Y | N.D. | N.D. | N.D. | N.D. | N.D. | N.D. | N.D. | N.D. | N.D. | N.D. | N.D. | N.D. | N.D. | N.D. |
| CAC231 | 14, 15 | 10, 12 | 11, 12 | 6 | 16, 19 | 28, 32.2 | 10, 11 | 11 | 8 | X, Y | N.D. | N.D. | N.D. | N.D. | N.D. | N.D. | N.D. | N.D. | N.D. | N.D. | N.D. | N.D. | N.D. | N.D. |
| CAC282 | 9, 12 | 8, 12 | 10 | 6, 7 | 14, 15 | 30 | N.D. | 11, 13 | 8, 11 | X, Y | N.D. | N.D. | N.D. | N.D. | N.D. | N.D. | N.D. | N.D. | N.D. | N.D. | N.D. | N.D. | N.D. | N.D. |
| SPCA | 11, 12 | 11 | 8, 11 | 8, 9.3 | 18 | 28, 30.2 | 10 | 11, 12 | 8 | X | N.D. | N.D. | N.D. | N.D. | N.D. | N.D. | N.D. | N.D. | N.D. | N.D. | N.D. | N.D. | N.D. | N.D. |
| WGS-CA (CA36) | 11, 12 | 11, 14 | 12 | 7, 10 | 16, 17 | 29, 31.2 | 10, 11 | 11, 13 | 8, 10 | X | N.D. | N.D. | N.D. | N.D. | N.D. | N.D. | N.D. | N.D. | N.D. | N.D. | N.D. | N.D. | N.D. | N.D. |
| JP-CA | 12 | 10, 11 | 10, 11 | 7 | 16, 18 | 31, 32.2 | N.D. | 12, 13 | 9 | X, Y | N.D. | N.D. | N.D. | N.D. | N.D. | N.D. | N.D. | N.D. | N.D. | N.D. | N.D. | N.D. | N.D. | N.D. |
| JW-CA | 12, 14 | 11, 13 | 12 | 6, 9.3 | 17 | 30, 32.2 | 9, 10 | 12 | 8, 10 | X | N.D. | N.D. | N.D. | N.D. | N.D. | N.D. | N.D. | N.D. | N.D. | N.D. | N.D. | N.D. | N.D. | N.D. |
| CRL7213 | 11, 12 | 12, 14 | 10, 12 | 8, 9 | 17, 18 | 28, 32.2 | 8, 14 | 11 | 8, 9 | N.D. | N.D. | N.D. | N.D. | N.D. | N.D. | N.D. | N.D. | N.D. | N.D. | N.D. | N.D. | N.D. | N.D. | N.D. |
| AACT (CT42) | 11, 13 | 11 | 11, 13 | 7, 9.3 | 18 | 31, 33.2 | 10 | 12 | 8, 9 | X | N.D. | N.D. | N.D. | N.D. | N.D. | N.D. | N.D. | N.D. | N.D. | N.D. | N.D. | N.D. | N.D. | N.D. |
| JW-NL | 12, 14 | 11, 13 | 12 | 6, 9.3 | 17 | 30, 32.2 | 9, 10 | 12 | 8, 10 | X | N.D. | N.D. | N.D. | N.D. | N.D. | N.D. | N.D. | N.D. | N.D. | N.D. | N.D. | N.D. | N.D. | N.D. |
| TVNL | N.D. | N.D. | N.D. | N.D. | N.D. | N.D. | N.D. | N.D. | N.D. | N.D. | N.D. | N.D. | N.D. | N.D. | N.D. | N.D. | N.D. | N.D. | N.D. | N.D. | N.D. | N.D. | N.D. | N.D. |
| NL322 | 13 | 9,11 | 9,11 | 6,7 | 16,18 | 27,30 | 11,12 | 12,13 | 8 | X,Y | N.D. | N.D. | N.D. | N.D. | N.D. | N.D. | N.D. | N.D. | N.D. | N.D. | N.D. | N.D. | N.D. | N.D. |
| PKNL (NL33) | 11 | 11,12 | 11,12 | 8, 9.3 | 15, 17 | 29, 32.2 | 10,12 | 9,11 | 8 | X,Y | N.D. | N.D. | N.D. | N.D. | N.D. | N.D. | N.D. | N.D. | N.D. | N.D. | N.D. | N.D. | N.D. | N.D. |
| CT5-SV40 | 11 | 9,11 | 12 | 6,7 | 16,17 | 29, 32.2 | 8 | 11,13 | 8,11 | X,Y | N.D. | N.D. | N.D. | N.D. | N.D. | N.D. | N.D. | N.D. | N.D. | N.D. | N.D. | N.D. | N.D. | N.D. |
| CRL1541-SV40 | 12 | 10, 11 | 9,13 | 6,7 | 16,18 | 28,30 | 8,9 | 12 | 8 | X | N.D. | N.D. | N.D. | N.D. | N.D. | N.D. | N.D. | N.D. | N.D. | N.D. | N.D. | N.D. | N.D. | N.D. |
| CT225-distal | 8, 12 | N.D. | 11, 14 | 6 | 16, 18 | 28, 30.2 | 10, 11 | 10, 12 | 8 | X, Y | N.D. | N.D. | N.D. | N.D. | N.D. | N.D. | N.D. | N.D. | N.D. | N.D. | N.D. | N.D. | N.D. | N.D. |
| CT225-proximal | 15, 18 | 12 | 11, 14 | 6 | 16, 18 | 28, 30.2 | 10, 11 | 10, 12 | 8 | X, Y | N.D. | N.D. | N.D. | N.D. | N.D. | N.D. | N.D. | N.D. | N.D. | N.D. | N.D. | N.D. | N.D. | N.D. |
| CT227-distal | 11,12 | 12,14 | 10,11 | 7,9.3 | 14,17 | 31 | 10 | 11,12 | 8,10 | X | N.D. | N.D. | N.D. | N.D. | N.D. | N.D. | N.D. | N.D. | N.D. | N.D. | N.D. | N.D. | N.D. | N.D. |
| CT227-proximal | 11,12 | 12,14 | 10,11 | 7,9.3 | 14,17 | 31 | 10 | 11,12 | 8,10 | X | N.D. | N.D. | N.D. | N.D. | N.D. | N.D. | N.D. | N.D. | N.D. | N.D. | N.D. | N.D. | N.D. | N.D. |
| CT279 distal | 17, 18 | 8, 12 | 10 | 6, 7 | 14, 15 | 30 | 11, 12 | 11, 13 | 8, 11 | X, Y | N.D. | N.D. | N.D. | N.D. | N.D. | N.D. | N.D. | N.D. | N.D. | N.D. | N.D. | N.D. | N.D. | N.D. |
| CT280-distal | N.D. | N.D. | N.D. | N.D. | N.D. | N.D. | N.D. | N.D. | N.D. | N.D. | N.D. | N.D. | N.D. | N.D. | N.D. | N.D. | N.D. | N.D. | N.D. | N.D. | N.D. | N.D. | N.D. | N.D. |
| CT281-distal | N.D. | N.D. | N.D. | N.D. | N.D. | N.D. | N.D. | N.D. | N.D. | N.D. | N.D. | N.D. | N.D. | N.D. | N.D. | N.D. | N.D. | N.D. | N.D. | N.D. | N.D. | N.D. | N.D. | N.D. |
| CT283-distal | N.D. | N.D. | N.D. | N.D. | N.D. | N.D. | N.D. | N.D. | N.D. | N.D. | N.D. | N.D. | N.D. | N.D. | N.D. | N.D. | N.D. | N.D. | N.D. | N.D. | N.D. | N.D. | N.D. | N.D. |
| CT284-distal | N.D. | N.D. | N.D. | N.D. | N.D. | N.D. | N.D. | N.D. | N.D. | N.D. | N.D. | N.D. | N.D. | N.D. | N.D. | N.D. | N.D. | N.D. | N.D. | N.D. | N.D. | N.D. | N.D. | N.D. |
| CT284-proximal | N.D. | N.D. | N.D. | N.D. | N.D. | N.D. | N.D. | N.D. | N.D. | N.D. | N.D. | N.D. | N.D. | N.D. | N.D. | N.D. | N.D. | N.D. | N.D. | N.D. | N.D. | N.D. | N.D. | N.D. |
| CT285-distal | 8, 12 | 11, 12 | 10, 11 | 7, 9 | 15,19 | 32.2 | 11, 12 | 11 | 11 | X,Y | N.D. | N.D. | N.D. | N.D. | N.D. | N.D. | N.D. | N.D. | N.D. | N.D. | N.D. | N.D. | N.D. | N.D. |
| CT285-proximal | 8,12 | 11, 12 | 10, 11 | 7, 9 | 15,19 | 32.2 | 11, 12 | 11 | 11 | X,Y | N.D. | N.D. | N.D. | N.D. | N.D. | N.D. | N.D. | N.D. | N.D. | N.D. | N.D. | N.D. | N.D. | N.D. |
| CT303-distal | 9,10 | 11,13 | 12 | 6,9.3 | 17,18 | 28,29 | 11,12 | 11, 12 | 8 | X,Y | N.D. | N.D. | N.D. | N.D. | N.D. | N.D. | N.D. | N.D. | N.D. | N.D. | N.D. | N.D. | N.D. | N.D. |
| CT303-proximal | 9,10 | 11,13 | 12 | 6,9.3 | 17,18 | 28,29 | 11,12 | 11, 12 | 8 | X,Y | N.D. | N.D. | N.D. | N.D. | N.D. | N.D. | N.D. | N.D. | N.D. | N.D. | N.D. | N.D. | N.D. | N.D. |
| CT333-distal | 11, 13 | 11, 13 | 10, 11 | 6 | 19, 20 | 29, 30 | N.D. | 11, 12 | 8, 11 | X, Y | N.D. | N.D. | N.D. | N.D. | N.D. | N.D. | N.D. | N.D. | N.D. | N.D. | N.D. | N.D. | N.D. | N.D. |
| CT333-proximal | 11, 13 | 11, 13 | 10, 11 | 6 | 19, 20 | 29, 30 | N.D. | 11, 12 | 8, 11 | X, Y | N.D. | N.D. | N.D. | N.D. | N.D. | N.D. | N.D. | N.D. | N.D. | N.D. | N.D. | N.D. | N.D. | N.D. |
| CT319-Prox | 12 | 9, 13 | 10, 11 | 6, 7 | 14, 17 | 28, 29 | 10, 11 | 10, 11 | 6, 7 | X, Y | N.D. | N.D. | N.D. | N.D. | N.D. | N.D. | N.D. | N.D. | N.D. | N.D. | N.D. | N.D. | N.D. | N.D. |
| CT319-distal | 12 | 9, 13 | 10, 11 | 6, 7 | 14, 17 | 28, 29 | 10, 11 | 10, 11 | 6, 7 | X, Y | N.D. | N.D. | N.D. | N.D. | N.D. | N.D. | N.D. | N.D. | N.D. | N.D. | N.D. | N.D. | N.D. | N.D. |
| CT320-distal | 8,9 | 9,11 | 10,12 | 8,9.3 | 16,18 | 27,30 | 8,10 | 10,13 | 9,11 | X | N.D. | N.D. | N.D. | N.D. | N.D. | N.D. | N.D. | N.D. | N.D. | N.D. | N.D. | N.D. | N.D. | N.D. |
| CT320-proximal | 8,9 | 9,11 | 10,12 | 8,9.3 | 16,18 | 27,30 | 8,10 | 10,13 | 9,11 | X | N.D. | N.D. | N.D. | N.D. | N.D. | N.D. | N.D. | N.D. | N.D. | N.D. | N.D. | N.D. | N.D. | N.D. |
| CT306-Proximal | N.D. | N.D. | N.D. | N.D. | N.D. | N.D. | N.D. | N.D. | N.D. | N.D. | N.D. | N.D. | N.D. | N.D. | N.D. | N.D. | N.D. | N.D. | N.D. | N.D. | N.D. | N.D. | N.D. | N.D. |
| CT306-distal-Tert | 12,13 | 12 | 12 | 8,9.3 | 16,18 | 28,30 | 9,10 | 11,13 | 8 | X | 11,16 | 11,14 | 18,25 | 15,17 | 13,15 | 13,14 | 20.3,24 | 13,17 | 14,15 | 15 |  | 21,22 | 13 | 7,12 |
| CT307-distal-Tert | 12,13 | 10,12 | 10,12 | 8,9.3 | 16,17 | 30,32.2 | 8,9 | 9,12 | 8,11 | X,Y | N.D. | N.D. | N.D. | N.D. | N.D. | N.D. | N.D. | N.D. | N.D. | N.D. | N.D. | N.D. | N.D. | N.D. |
| CT307-proximal-Tert | 12,13 | 10,12 | 10,12 | 8,9.3 | 16,17 | 30,32.2 | 8,9 | 9,12 | 8,11 | X,Y | N.D. | N.D. | N.D. | N.D. | N.D. | N.D. | N.D. | N.D. | N.D. | N.D. | N.D. | N.D. | N.D. | N.D. |
| CT350 prox | 9,11 | 12,13 | 11 | 7,8 | 14,16 | 28,29 | 8,11 | 12,13 | 7,9 | X,Y | 11,18.3 | 11,14 | 16,17 | 16,18 | 12,13 | 13,14 | 20 | 18,21 | 11,15 | 15 | 11 | 19,22 | 2.2,12 | 9,14 |
| CT350 dist | 9,11 | 12,13 | 11 | 7,8 | 14,16 | 28,29 | 8,11 | 12,13 | 7,9 | X,Y | 11,18.3 | 11,14 | 16,17 | 16,18 | 12,13 | 13,14 | 20 | 18,21 | 11,15 | 15 | 11 | 19,22 | 2.2,12 | 9,14 |
| CT351 Prox | 9,12 | 11,12 | 12 | 6,9 | 14,19 | 30,32.2 | 11,13 | 11 | 8,10 | X | 13,15 | 10 | 17 | 17,18 | 10,12 | 15,16 | 21,22 | 14,17 | 12,16.2 | 15 | N.D. | 23 | 9 | 7,13 |
| CT351 dist | 9,12 | 11,12 | 12 | 6,9 | 14,19 | 30,32.2 | 11,13 | 11 | 8,10 | X | 13,15 | 10 | 17 | 17,18 | 10,12 | 15,16 | 21,22 | 14,17 | 12,16.2 | 15 | N.D. | 23 | 9 | 7,13 |
| HCT-116 (CCL-247) | 10,12,13 | 11,13 | 7,10 | 8,9 | 16, 17,22 | 29,30 | 11,12 | 10,11 | 8 | X | N.D. | N.D. | N.D. | N.D. | N.D. | N.D. | N.D. | N.D. | N.D. | N.D. | N.D. | N.D. | N.D. | N.D. |
| CT420C Prox | N.D. | N.D. | N.D. | N.D. | N.D. | N.D. | N.D. | N.D. | N.D. | N.D. | In Dewar 3 | C4 box 7 | N.D. | N.D. | N.D. | N.D. | N.D. | N.D. | N.D. | N.D. | N.D. | N.D. | N.D. | N.D. |
| CT420C Distal | N.D. | N.D. | N.D. | N.D. | N.D. | N.D. | N.D. | N.D. | N.D. | N.D. | N.D. | N.D. | N.D. | N.D. | N.D. | N.D. | N.D. | N.D. | N.D. | N.D. | N.D. | N.D. | N.D. | N.D. |
| CA404C | 11 | 10, 11 | 10 | 6 | 14,16 | 29, 30 | 9, 11 | 13 | 8,10 | X | N.D. | N.D. | N.D. | N.D. | N.D. | N.D. | N.D. | N.D. | N.D. | N.D. | N.D. | N.D. | N.D. | N.D. |
| CT461-Proximal | 12 | 11,12 | 10,12 | 9,9.3 | 18 | 32.2,33.2 | 11 | 9 | 8,9 | X | N.D. | N.D. | N.D. | N.D. | N.D. | N.D. | N.D. | N.D. | N.D. | N.D. | N.D. | N.D. | N.D. | N.D. |
| CT461-Distal | 12 | 11,12 | 10,12 | 9,9.3 | 18 | 32.2,33.2 | 11 | 9 | 8,9 | X | N.D. | N.D. | N.D. | N.D. | N.D. | N.D. | N.D. | N.D. | N.D. | N.D. | N.D. | N.D. | N.D. | N.D. |

Supplementary Table S2: STR Analysis
